# Genetic Determinants of Cortical Structure (Thickness, Surface Area and Volumes) among Disease Free Adults in the CHARGE Consortium

**DOI:** 10.1101/409649

**Authors:** Edith Hofer, Gennady V. Roshchupkin, Hieab H. H. Adams, Maria J. Knol, Honghuang Lin, Shuo Li, Habil Zare, Shahzad Ahmad, Nicola J. Armstrong, Claudia L. Satizabal, Manon Bernard, Joshua C. Bis, Nathan A. Gillespie, Michelle Luciano, Aniket Mishra, Markus Scholz, Alexander Teumer, Rui Xia, Xueqiu Jian, Thomas H. Mosley, Yasaman Saba, Lukas Pirpamer, Stephan Seiler, James T. Becker, Owen Carmichael, Jerome I. Rotter, Bruce M. Psaty, Oscar L. Lopez, Najaf Amin, Sven J. van der Lee, Qiong Yang, Jayandra J. Himali, Pauline Maillard, Alexa S. Beiser, Charles DeCarli, Sherif Karama, Lindsay Lewis, Mat Harris, Mark E. Bastin, Ian J. Deary, A.Veronica Witte, Frauke Beyer, Markus Loeffler, Karen A. Mather, Peter R. Schofield, Anbupalam Thalamuthu, John B. Kwok, Margaret J. Wright, David Ames, Julian Trollor, Jiyang Jiang, Henry Brodaty, Wei Wen, Meike W Vernooij, Albert Hofman, André G. Uitterlinden, Wiro J. Niessen, Katharina Wittfeld, Robin Bülow, Uwe Völker, Zdenka Pausova, G. Bruce Pike, Sophie Maingault, Fabrice Crivello, Christophe Tzourio, Philippe Amouye, Bernard Mazoyer, Michael C. Neale, Carol E. Franz, Michael J. Lyons, Matthew S. Panizzon, Ole A. Andreassen, Anders M. Dale, Mark Logue, Katrina L. Grasby, Neda Jahanshad, Jodie N. Painter, Lucía Colodro-Conde, Janita Bralten, Derrek P. Hibar, Penelope A. Lind, Fabrizio Pizzagalli, Jason L. Stein, Paul M. Thompson, Sarah E. Medland

## Abstract

Cortical thickness, surface area and volumes (MRI cortical measures) vary with age and cognitive function, and in neurological and psychiatric diseases. We examined heritability, genetic correlations and genome-wide associations of cortical measures across the whole cortex, and in 34 anatomically predefined regions. Our discovery sample comprised 22,824 individuals from 20 cohorts within the Cohorts for Heart and Aging Research in Genomic Epidemiology (CHARGE) consortium and the United Kingdom Biobank. Significant associations were replicated in the Enhancing Neuroimaging Genetics through Meta-analysis (ENIGMA) consortium, and their biological implications explored using bioinformatic annotation and pathway analyses. We identified genetic heterogeneity between cortical measures and brain regions, and 160 genome-wide significant associations pointing to wnt/β-catenin, TGF-β and sonic hedgehog pathways. There was enrichment for genes involved in anthropometric traits, hindbrain development, vascular and neurodegenerative disease and psychiatric conditions. These data are a rich resource for studies of the biological mechanisms behind cortical development and aging.

## Introduction

The cortex is the largest part of the human brain, associated with higher brain functions such as perception, thought and action. Brain cortical thickness (CTh), cortical surface area (CSA) and cortical volume (CV) are morphological markers of cortical structure obtained from magnetic resonance imaging (MRI). These measures change with age^1-3^ and are linked to cognitive functioning^4,5^. The human cortex is also vulnerable to a wide range of disease or pathologies, ranging from developmental disorders and early onset psychiatric and neurological diseases to neurodegenerative conditions manifesting late in life. Abnormalities in global or regional CTh, CSA and CV have been observed in neurological and psychiatric disorders such as Alzheimer’s disease^6,7^, Parkinson’s disease^8,9^, multiple sclerosis^10,11^, schizophrenia^12,13^, bipolar disorder^12,14,15^, depression^15,16^ and autism^17,18^. The best method to study human cortical structure during life is using brain MRI. Hence, understanding the genetic determinants of the most robust MRI cortical markers in apparently normal adults could identify biological pathways relevant to brain development, aging and various diseases. Neurons in the neocortex are organized in columns which run perpendicular to the surface of the cerebral cortex^19^; and, according to the radial unit hypothesis, CTh is determined by the number of cells within the columns and CSA is determined by the number of columns^20^. Thus, CTh and CSA reflect different mechanisms in cortical development^20-24^ and are likely influenced by different genetic factors^25,26^. CV, which is the product of CTh and CSA, is determined by a combination of these two measures, but the relative contribution of CTh and CSA to CV may vary across brain regions. CTh, CSA and CV are all strongly heritable traits^22,24-30^ with estimated heritability of 0.69 to 0.81 for global CTh, and from 0.42 to 0.90 for global CSA^24-26^. Across different cortical regions however, there is substantial regional variation in heritability of CTh, CSA and CV^22,24-30^. Since CTh, CSA and CV are differentially heritable and genetically heterogeneous, we explored the genetics of each ofthese imaging markers using genome-wide association analyses in large population-based samples (GWAS). We studied CTh, CSA and CV in the whole cortex and in 34 cortical regions in 22,824 individuals from 21 discovery cohorts and replicated the strongest associations in 22,363 persons from the ENIGMA consortium.

## Results

### Genome-wide association analysis

#### Global Cortical Measures

The analyses of global CTh, CSA and CV included 22,163, 18,617 and 22,824 individuals respectively. After a conservative correction for multiple testing (p_discovery_<1.09×10^-9^), we identified no significant associations with global CTh. However, we identified 12 independent loci associated with global CSA (n=6) and CV (n=6). These are displayed in Table S8 and Supplementary Figures 1 and 2. Five of the 6 CSA loci were replicated in an external (ENIGMA consortium) sample^31^. The ENIGMA consortium only analyzed CSA and CTh.

#### Regional Cortical Measures

GWAS of CTh, CSA and CV in 34 cortical regions of interest (ROIs) identified 148 significant associations. There were 16 independent loci across 8 chromosomes determining CTh of 9 regions (Table S9), 54 loci across 16 chromosomes associated with CSA of 21 regions (Table S10), and 78 loci across 17 chromosomes determining CV of 23 cortical regions (Table S11). We replicated 57 out of 70 regional CTh and CSA loci in the ENIGMA consortium sample^31^ using a conservative replication threshold of p_Replication_=3.1×10^-4^, 0.05/160. Region-specific variants with the strongest association at each genomic locus are shown in Tables 1-3. Chromosomal ideograms showing genome-wide significant associations with global and regional cortical measures in the discovery stage are presented in Figure 1.

**Figure 1.**
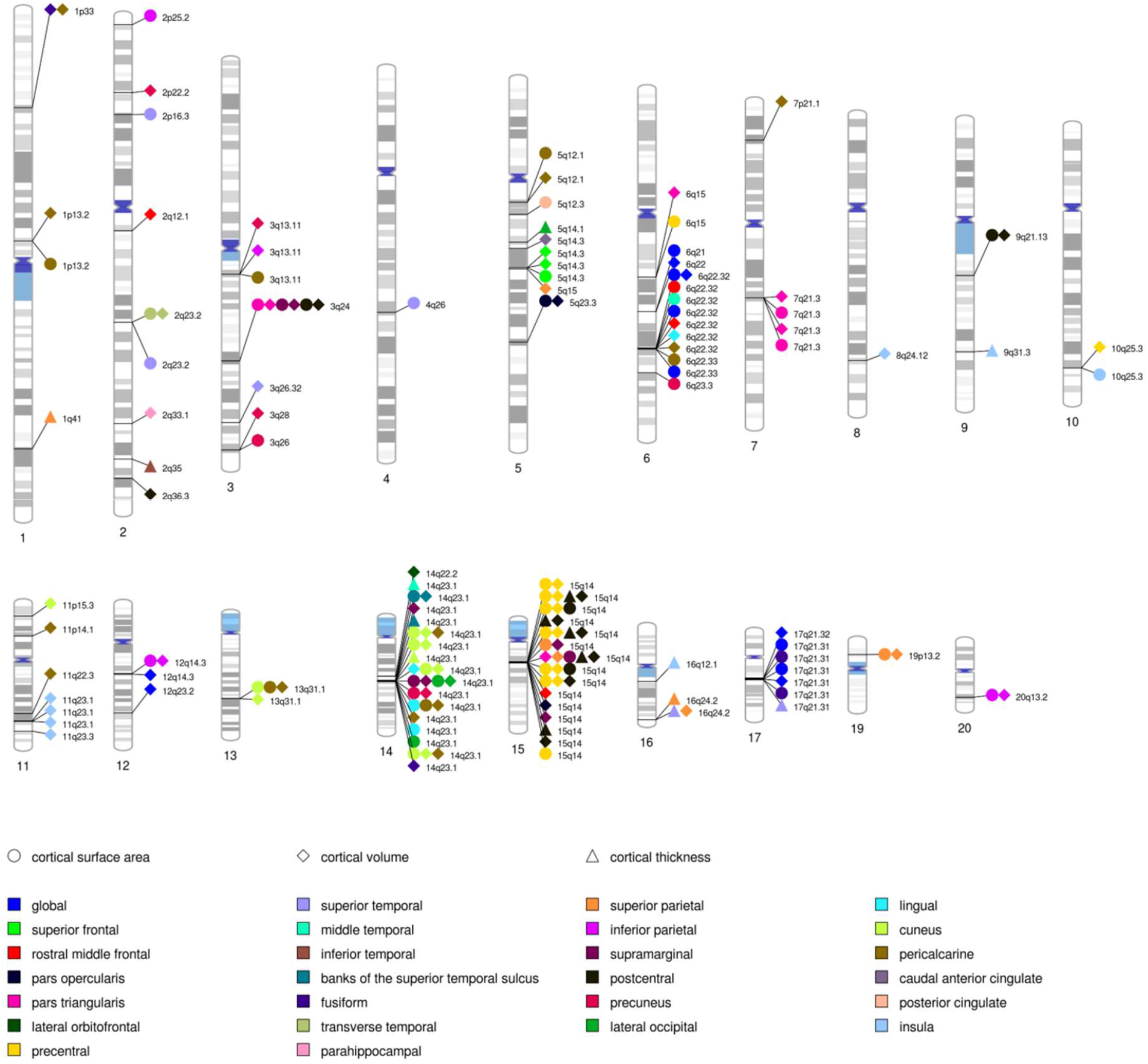
Chromosomal ideogram annotated with genome-wide significant associations (p_Discovery_<1.09×10^-9^) and corresponding genomic loci.

**Table 1.**
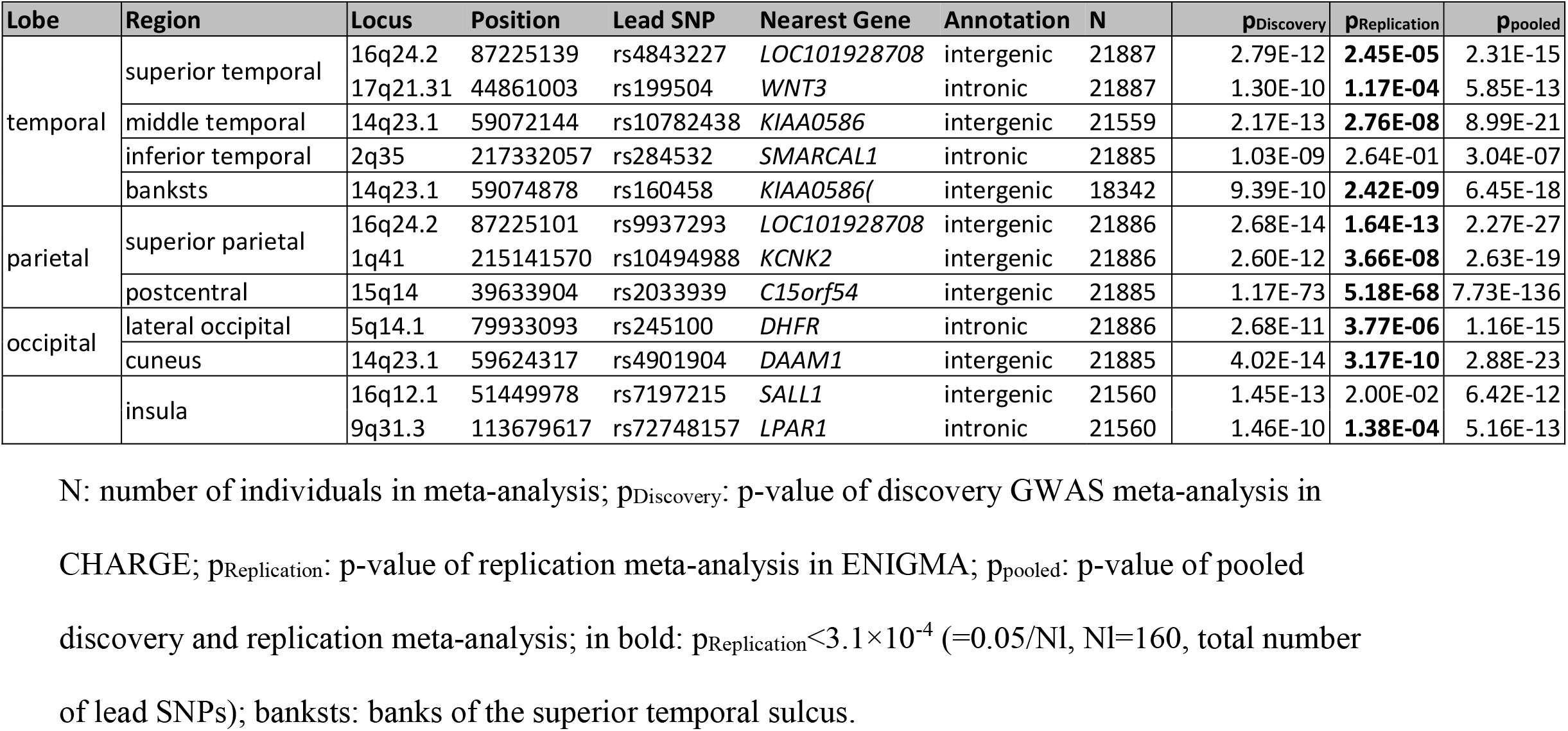
Genome-wide significant associations (p_Discovery_ < 1.09×10-9) of regional cortical thickness (lowest p-value of each cortical region at each genomic locus)

**Table 2.**
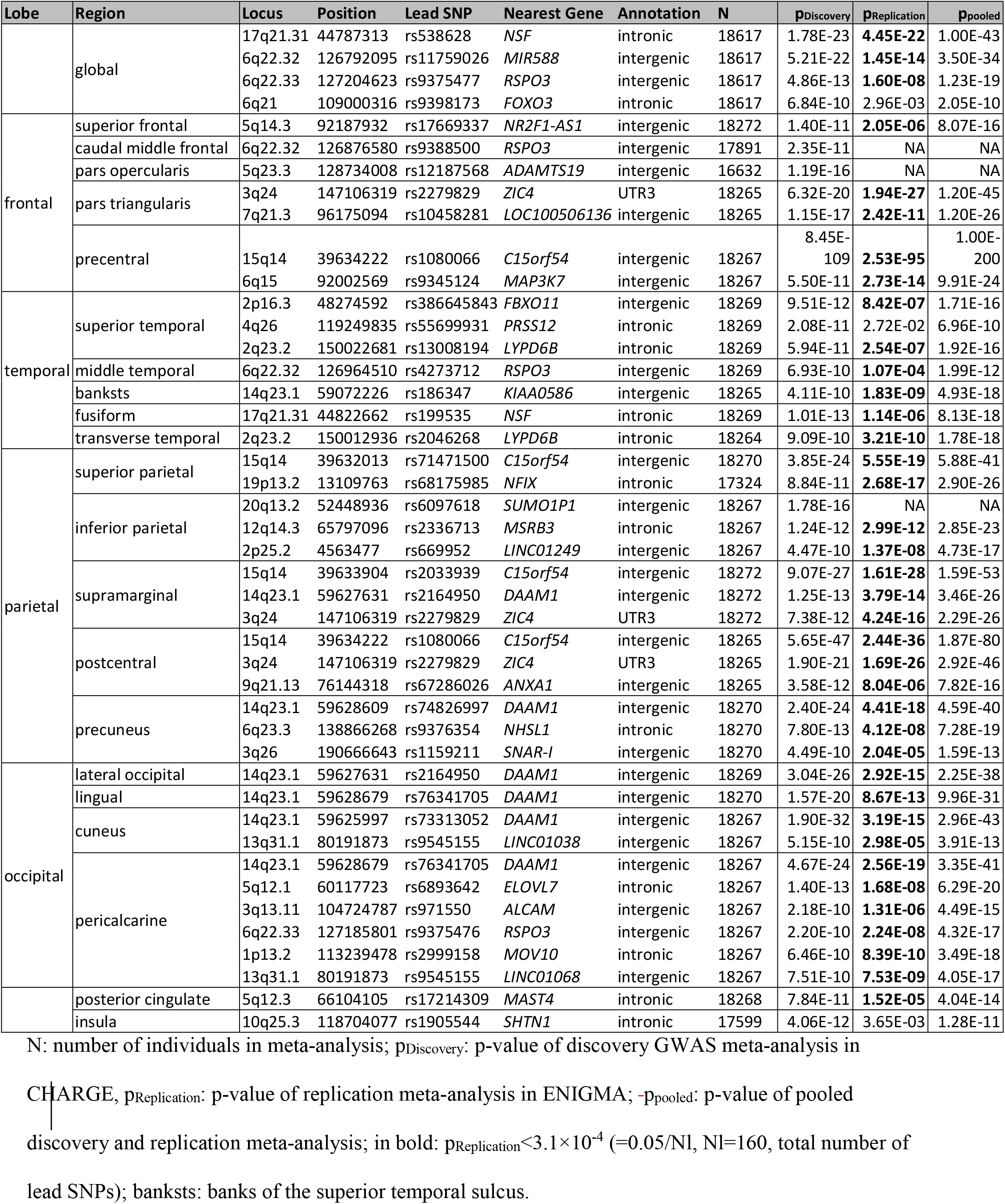
Genome-wide significant associations (p_Discovery_ < 1.09×10-9) of global and regional cortical surface area (lowest p-value of each cortical region at each genomic locus)

**Table 3.** Genome-wide significant associations (p_Discovery_ < 1.09×10-9) of global and regional cortical volume (lowest p-value of each cortical region at each genomic locus)

| Lobe | Region | Locus | Position | Lead SNP | Nearest gene | Annotation | N | $p_{\text{Discovery}}$ |
| --- | --- | --- | --- | --- | --- | --- | --- | --- |
|  | global | 6q22.32 | 126792095 | rs11759026 | *MIR588* | intergenic | 22410 | 6.31E-19 |
|  |  | 17q21.31 | 44790203 | rs169201 | *NSF* | intronic | 22784 | 2.11E-13 |
|  |  | 17q21.32 | 43549608 | rs149366495 | *PLEKHM1* | intronic | 22099 | 8.18E-13 |
|  |  | 12q14.3 | 66358347 | rs1042725 | *HMGA2* | 3'UTR | 22784 | 7.04E-11 |
|  |  | 12q23.2 | 102921296 | rs11111293 | *IGF1* | intergenic | 22784 | 5.45E-10 |
|  |  | 6q22 | 109002042 | rs4945816 | *FOXO3* | 3'UTR | 22784 | 8.93E-10 |
| frontal | superior frontal | 5q14.3 | 92186429 | rs888814 | *NR2F1-AS1* | intergenic | 22692 | 3.29E-13 |
|  | rostral middle frontal | 15q14 | 39636227 | rs17694988 | *C15orf54* | intergenic | 22793 | 3.15E-11 |
|  | caudal middle frontal | 2q12.1 | 105460333 | rs745249 | *LINC01158* | ncRNA_intronic | 22726 | 2.35E-11 |
|  |  | 6q22.32 | 127068983 | rs853974 | *RSPO3* | intergenic | 22351 | 4.82E-11 |
|  | pars opercularis | 5q23.3 | 128734008 | rs12187568 | *ADAMTS19* | intergenic | 20753 | 4.27E-18 |
|  |  | 15q14 | 39639898 | rs4924345 | *C15orf54* | intergenic | 22758 | 1.97E-14 |
|  | pars triangularis | 3q24 | 147106319 | rs2279829 | *ZIC4* | UTR3 | 22759 | 3.16E-23 |
|  |  | 7q21.3 | 96196906 | rs67055449 | *LOC100506136* | intergenic | 22759 | 4.03E-19 |
|  |  | 15q14 | 39633904 | rs2033939 | *C15orf54* | intergenic | 22759 | 8.49E-14 |
|  |  | 7q21.3 | 96129071 | rs62470042 | *C7orf76* | intronic | 22759 | 7.38E-13 |
|  |  | 6q15 | 91942761 | rs12660096 | *MAP3K7* | intergenic | 22759 | 4.74E-10 |
|  | lateral orbitofrontal | 14q22.2 | 54769839 | rs6572946 | *CDKN3* | intergenic | 22801 | 2.29E-10 |
|  | precentral | 15q14 | 39634222 | rs1080066 | *C15orf54* | intergenic | 22699 | 5.84E-125 |
|  |  | 10q25.3 | 118648841 | rs3781566 | *SHTN1* | intronic | 22699 | 4.68E-11 |
| temporal | superior temporal | 3q26.32 | 177296448 | rs13084960 | *LINC00578* | ncRNA_intronic | 22681 | 1.12E-11 |
|  | banksts | 14q23.1 | 59072226 | rs186347 | *KIAA0586* | intergenic | 22727 | 1.15E-15 |
|  | fusiform | 14q23.1 | 59833172 | rs1547199 | *DAAM1* | intronic | 22605 | 4.58E-10 |
|  |  | 1p33 | 47980916 | rs6658111 | *FOXD2* | intergenic | 22605 | 7.78E-10 |
|  | transverse temporal | 2q23.2 | 150012936 | rs2046268 | *LYPD6B* | intronic | 22786 | 2.55E-12 |
|  | parahippocampal | 2q33.1 | 199809716 | rs966744 | *SATB2* | intergenic | 22747 | 2.23E-10 |
| parietal | superior parietal | 15q14 | 39633904 | rs2033939 | *C15orf54* | intergenic | 22723 | 4.28E-23 |
|  |  | 16q24.2 | 87225139 | rs4843227 | *LOC101928708* | intergenic | 22723 | 1.16E-13 |
|  |  | 19p13.2 | 13109763 | rs68175985 | *NFIX* | intronic | 21777 | 3.27E-11 |
|  |  | 5q15 | 92866553 | rs62369942 | *NR2F1-AS1* | ncRNA_intronic | 21664 | 4.32E-10 |
|  | inferior parietal | 20q13.2 | 52448936 | rs6097618 | *SUMO1P1* | intergenic | 22701 | 2.09E-17 |
|  |  | 12q14.3 | 65797096 | rs2336713 | *MSRB3* | intronic | 22701 | 2.47E-13 |
|  |  | 3q13.11 | 104724634 | rs971551 | *ALCAM* | intergenic | 22701 | 2.34E-10 |
|  | supramarginal | 15q14 | 39632013 | rs71471500 | *THBS1* | intergenic | 22645 | 9.71E-28 |
|  |  | 14q23.1 | 59627631 | rs2164950 | *DAAM1* | intergenic | 22645 | 3.59E-20 |
|  |  | 3q24 | 147106319 | rs2279829 | *ZIC4* | UTR3 | 22645 | 5.36E-18 |
|  | postcentral | 15q14 | 39633904 | rs2033939 | *THBS1* | intergenic | 22662 | 4.34E-133 |
|  |  | 3q24 | 147106319 | rs2279829 | *ZIC4* | UTR3 | 22662 | 2.54E-17 |
|  |  | 9q21.13 | 76144318 | rs67286026 | *ANXA1* | intergenic | 22662 | 5.03E-11 |
|  |  | 2q36.3 | 226563259 | rs16866701 | *NYAP2* | intergenic | 22545 | 5.69E-11 |
|  | precuneus | 14q23.1 | 59628609 | rs74826997 | *DAAM1* | intergenic | 22803 | 4.85E-20 |
|  |  | 3q28 | 190663557 | rs35055419 | *OSTN* | intergenic | 22428 | 2.02E-10 |
|  |  | 2p22.2 | 37818236 | rs2215605 | *CDC42EP3* | intergenic | 22803 | 3.43E-10 |
|  |  | 3q13.11 | 104713881 | rs12495603 | *ALCAM* | intergenic | 22803 | 9.71E-10 |
| occipital | lateral occipital | 14q23.1 | 59627631 | rs2164950 | *DAAM1* | intergenic | 22799 | 6.89E-16 |
|  | lingual | 14q23.1 | 59625997 | rs73313052 | *DAAM1* | intergenic | 22805 | 1.06E-20 |
|  |  | 6q22.32 | 127089401 | rs2223739 | *RSPO3* | intergenic | 22805 | 1.75E-10 |
|  | cuneus | 14q23.1 | 59625997 | rs73313052 | *DAAM1* | intergenic | 22799 | 4.59E-43 |
|  |  | 11p15.3 | 12072213 | rs11022131 | *DKK3* | intergenic | 22799 | 5.96E-12 |
|  |  | 13q31.1 | 80192236 | rs9545156 | *LINC01068* | intergenic | 22799 | 4.09E-10 |
|  | pericalcarine | 14q23.1 | 59628679 | rs76341705 | *DAAM1* | intergenic | 22824 | 1.39E-29 |
|  |  | 13q31.1 | 80191873 | rs9545155 | *LINC01068* | intergenic | 22824 | 2.25E-13 |
|  |  | 11p14.1 | 30876113 | rs273594 | *DCDC5* | intergenic | 22824 | 3.51E-13 |
|  |  | 1p13.2 | 113208039 | rs12046466 | *CAPZA1* | intronic | 22824 | 2.36E-12 |
|  |  | 1p33 | 47980916 | rs6658111 | *FOXD2* | intergenic | 22824 | 3.85E-11 |
|  |  | 11q22.3 | 104012656 | rs1681464 | *PDGFD* | intronic | 22824 | 7.51E-11 |
|  |  | 6q22.32 | 127096181 | rs9401907 | *RSPO3* | intergenic | 22824 | 2.11E-10 |
|  |  | 7p21.1 | 18904400 | rs12700001 | *HDAC9* | intronic | 22824 | 2.12E-10 |
|  |  | 5q12.1 | 60315823 | rs10939879 | *NDUFAF2* | intronic | 22824 | 2.92E-10 |
|  | caudal anterior cingulate | 5q14.3 | 82852578 | rs309588 | *VCAN* | intronic | 22748 | 2.60E-10 |
|  | insula | 11q23.1 | 110949402 | rs321403 | *C11orf53* | intergenic | 22543 | 9.58E-12 |
|  |  | 8q24.12 | 120596023 | rs10283100 | *ENPP2* | exonic | 21481 | 8.29E-11 |

N: number of individuals in meta-analysis; $p_{\text{Discovery}}$ : p-value of discovery GWAS meta-analysis in
CHARGE; banksts: banks of the superior temporal sulcus.

The strongest associations with CTh and CV were observed for rs2033939 at 15q14 (p_Discovery, CTh_=1.17×10^-73^ and p_Discovery, CV_=4.34×10^-133^) in the postcentral (primary somatosensory) cortex, and for CSA with rs1080066 at 15q14 (p_Discovery, CSA_=8.45×10^-109^) in the precentral (primary motor) cortex. Figure 2 shows the lowest p-value of each cortical region. The postcentral cortex was also the region with the largest number of independent associations, mainly at a locus on 15q14. The corresponding regional association plots are presented in Supplementary Figure 3.

QQ plots of all meta-analyses are presented in Supplementary Figures 4-7.

**Figure 2.**
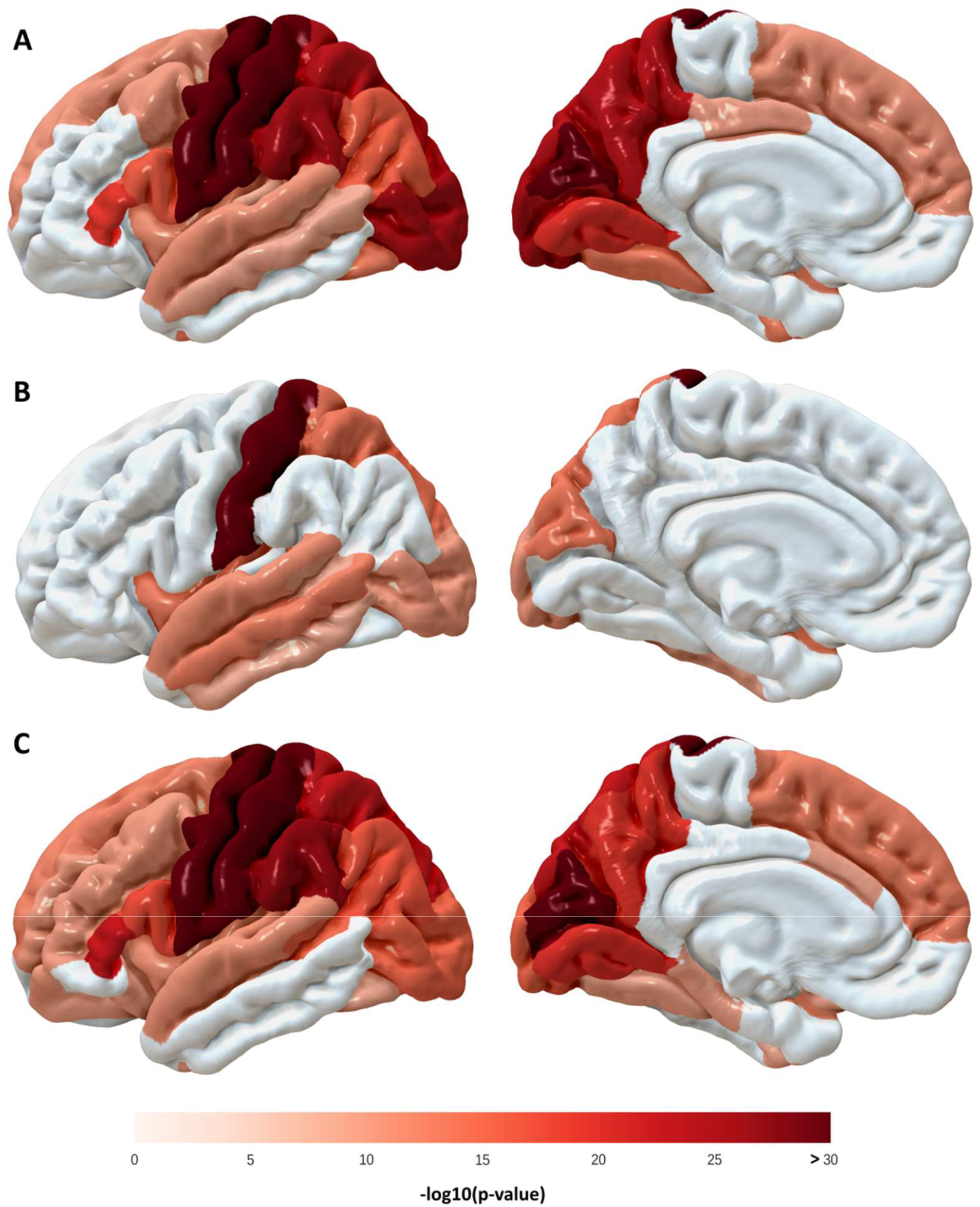
Lowest p-value of cortical surface area (A), thickness (B) and (C) volume of each cortical region.

#### Associations across Cortical Measures and with Other Traits

Table S12 presents variants which are associated with the CSA or the CV across multiple regions. We observed 25 SNPs that determined both the CSA and CV of a given region, 4 SNPs that determined CTh and CV of the same region, but no SNPs that determined both the CTh and CSA of any given region (Table S13). In the UK Biobank^32^, 3 SNPs were associated with the same cortical measure at a given cortical region as in our study. These included associations between rs2033939 at 15q14 and postcentral CTh, rs2279829 at 3q24 and CSA of the pars triangularis and postcentral gyrus and rs76341705 at 14q23.1 and CSA of lingual gyrus, respectively (Table S14). When assessing genetic overlap with other traits, we observed that SNPs determining these cortical measures have been previously associated with anthropometric (height), neurologic (Parkinson’s disease, corticobasal degeneration, Alzheimer’s disease), psychiatric (neuroticism, schizophrenia) and cognitive performance traits as well as with total intracranial volume (TIV) on brain MRI (Tables S15-S17).

### Gene Identification

Positional mapping based on ANNOVAR showed that most of the lead SNPs were intergenic and intronic (Figure 3). One variant, rs2279829, which was associated with both CSA and CV of the pars triangularis, postcentral and supramarginal cortices, is located in the 3’prime UTR of *ZIC4* at 3q24. We also found an exonic variant, rs10283100, in gene *ENPP2* at 8q24.12 associated with CV of the insula.

**Figure 3.**
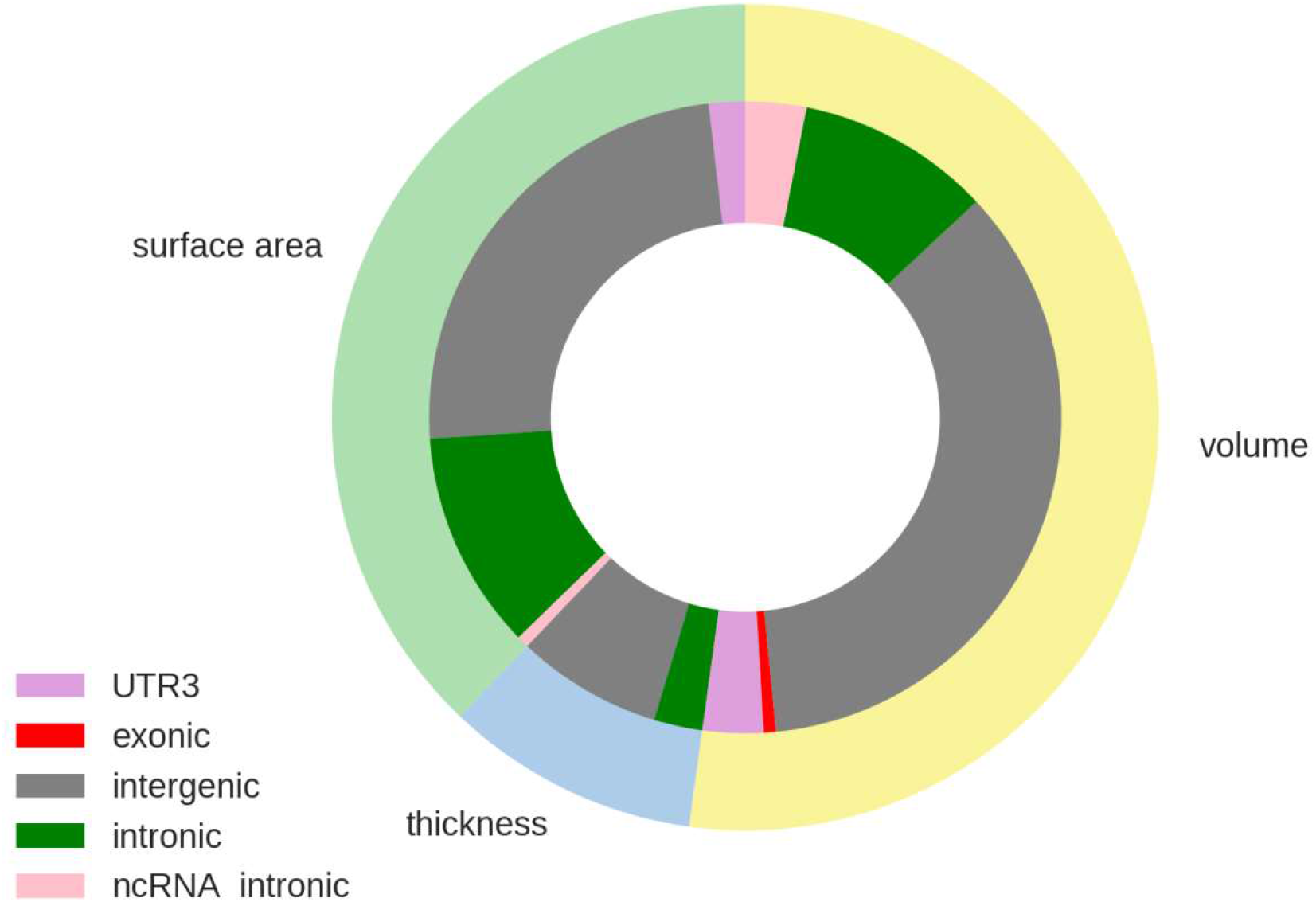
Proportion of functional annotation categories for global and regional cortical thickness, surface area and volume assigned by ANNOVAR.

We used multiple strategies beyond positional annotation to identify specific genes implicated by the various GWAS associated SNPs. FUMA identified 232 genes whose expression was determined by these variants (eQTL) and these and other genes implicated by chromatin interaction mapping are shown in Tables S18 – S20. MAGMA gene-based association analyses revealed 70 significantly associated (p<5.87*10^-8^) genes (Tables S21 – S23). For global CSA and CV, 7 of 9 genes associated with each measure overlapped, but there was no overlap with global CTh. For regional CSA and CV we found 28 genes across 13 cortical regions that determined both measures in the same region. Figure 4 summarizes the results of GTEx eQTL, chromatin interaction, positional annotation and gene-based mapping strategies for all regions. While there are overlapping genes identified using different approaches, only *DAAM1* gene (Chr14q23.1) is identified by all types of gene mapping for CV of insula. eQTL associations of our independent lead SNPs in the Religious Orders Study- Memory and Aging Project dorsolateral frontal cortex gene expression dataset are presented in Table S24.

**Figure 4.**
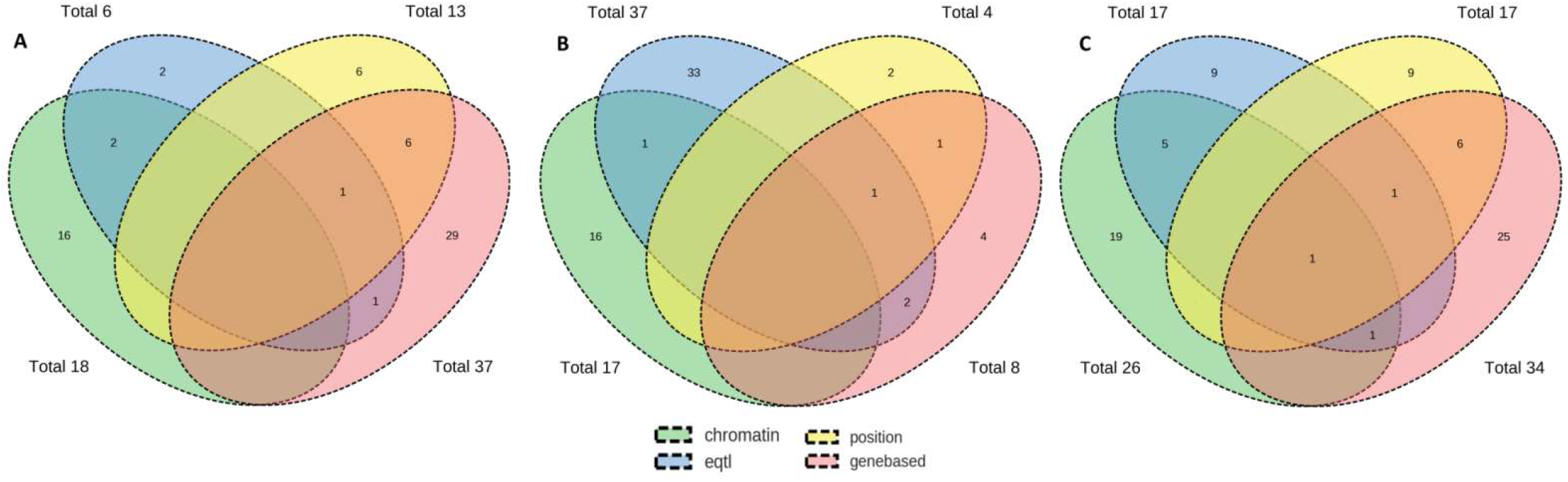
Number of overlapping genes between FUMA eQTL mapping, FUMA chromatin interaction mapping, ANNOVAR chromosome positional mapping and MAGMA gene based analysis for all cortical regions combined for cortical surface area (A), thickness (B) and volume (C).

### Pathway analysis

MAGMA gene set analyses identified 7 pathways for CTh, 3 pathways for CSA and 9 pathways for CV (Table S25). Among them are the Gene Ontology (GO) gene sets ‘hindbrain morphogenesis’ (strongest association with thickness of middle temporal cortex), ‘forebrain generation of neurons’ (with surface area of precentral cortex), and ‘central nervous system neuron development’ (with volume of transverse temporal cortex). However, after Bonferroni correction only one significant pathway (p<1.02×10^-7^) remained: ‘regulation of catabolic process’ for CTh of inferior temporal cortex. InnateDB pathway analyses of genes mapped to independent lead SNPs by FUMA showed a significant overlap between CTh and CSA genes and the Wnt signaling pathway (Supplementary Figures 8 and 9) as well as a significant overlap between CV genes and the basal cell carcinoma pathway (Supplementary Figure 10).

### Heritability

Heritability estimates (h^2^) of global CTh were 0.64 (se=0.12; p=3×10^-7^) in ASPS-Fam and 0.45 (se=0.08; p=2.5×10^-7^) in RS. For CSA, h^2^ was 0.84 (se=0.12; p=2.63×10^-11^) in ASPS-Fam and 0.33 (se=0.08, p=1×10^-4^) in RS, and for CV, h^2^ was 0.80 (se=0.11; p=1.10×10^-9^) in ASPS-Fam and 0.32 (se=0.08; p=1×10^-4^) in RS. There was a large range in heritability estimates of regional CTh, CSA and CV (Table S26).

Heritability based on common SNPs as estimated with LDSR was 0.25 (se=0.03) for global CTh, 0.29 (se=0.04) for global CSA and 0.30 (se=0.03) for global CV. LDSR heritability estimates of regional CTh, CSA and CV are presented in Table S26 and Supplementary Figure 11. For the regional analyses, the estimated heritability ranged from 0.05 to 0.18 for CTh, from 0.07 to 0.36 for CSA and from 0.06 to 0.32 for CV. Superior temporal cortex (h^2^_CTh_=0.18, h^2^_CSA_ =0.30, h^2^_cv_ =0.26), precuneus (h^2^_cTh_ =0.16, h^2^_cSA_ =0.29, h^2^_CV_ =0.28) and pericalcarine (h^2^_CTh_=0.15, h^2^_CSA_=0.36, h^2^_CV_=0.32) are among the most genetically determined regions.

The results of partitioned heritability analyses for global and regional CTh, CSA and CV with functional annotation and additionally with cell-type specific annotation are presented in Tables S27 and S28. For global CTh we found enrichment for super-enhancers, introns and histone marks. Repressors and histone marks were enriched for global CSA, and introns, super-enhancers and repressors for global CV. For regional CSA and CV the highest enrichment scores (>18) were observed for conserved regions.

### Genetic correlation

We found high genetic correlation (r_g_) between global CSA and global CV (r_g_=0.81, p=1.2×10^-186^) and between global CTh and global CV (r_g_=0.46, p=1.4×10^-14^), but not between global CTh and global CSA (r_g_= −0.02, p=0.82). Whereas genetic correlation between CSA and CV was strong (r_g_ >0.7) in most of the regions (Table S29 and Supplementary Figure 12), it was generally weak between CSA and CTh with r_g_<0.3, and ranged from 0.09 to 0.69 between CTh and CV. The postcentral and lingual cortices were the two regions with the highest genetic correlations between both CTh and CV, as well as CTh and CSA.

Genetic correlation across the various brain regions for CTh (Supplementary Figure 13, Table S30), CSA (Supplementary Figure 14, Table S30), and CV (Supplementary Figure 15, Table S32) showed a greater number of correlated regions for CTh and greater inter-regional variation for CSA and CV. Tables S33 - S35 and Supplementary Figures 16-18 show genome-wide genetic correlations between the cortical measures and anthropometric, neurological and psychiatric, and cerebral structural traits.

## Discussion

In our genome-wide association study of up to 22,824 individuals for MRI determined cortical measures of global and regional thickness, surface area and volume, we identified 160 genome-wide significant associations across 19 chromosomes. Heritability was generally higher for cortical surface area and volume than for thickness, suggesting a greater susceptibility of cortical thickness to environmental influences. We observed strong genetic correlations between surface area and volume, but weak genetic correlation between surface area and thickness. We identified the largest number of novel genetic associations with cortical volumes, perhaps due to our larger sample size for this phenotype which was assessed in all 21 discovery samples.

It is beyond the scope of our study to discuss each of the 160 associations identified. However, broad patterns emerged showing that genes determining cortical structure are also often implicated in development of the cerebellum and brainstem (*KIAA0586, ZIC4, ENPP2*) as well as the neural tube (one carbon metabolism genes *DHFR* and *MSRBB3,* the latter also associated with hippocampal volumes^33^). These genes determine development of not only neurons but also astroglia (*THBS1*) and microglia (*SALL1*). They determine susceptibility or resistance to a range of insults: inflammatory, vascular (*THBS1, ANXA1, ARRDC3-AS1*^*34*^) and neurodegenerative (*C15orf53, ZIC4, ANXA1*), and have been associated with pediatric and adult psychiatric conditions (*THBS1*). At a molecular level, the wnt/β-catenin, TGF-β and sonic hedgehog pathways are strongly implicated. Gene-set-enrichment analyses revealed biological processes related to brain morphology and neuronal development.

There is a wealth of information in the supplementary tables that can be mined for a better understanding of brain development, connectivity, function and pathology. We highlight this potential by discussing in additional detail, the possible significance of 6 illustrative loci, 5 of which, at 15q14, 14q23.1, 6q22.32, 17q21.31 and 3q24, associate with multiple brain regions at low p-values, while the locus at 8q24.12 identifies a plausible exonic variant.

The Chr15q14 locus was associated with cortical thickness, surface area and volumes in the postcentral gyrus as well as with surface area or volume across 6 other regions in the frontal and parietal lobes. Lead SNPs at this locus were either intergenic between *C15orf53* and *C15orf54*, or intergenic between *C15orf54* and *THBS1* (Thrombospondin-1). *C15orf53* has been associated with an autosomal recessive form of spastic paraplegia showing intellectual disability and thinning of the corpus callosum (hereditary spastic paraparesis 11, or Nakamura Osame syndrome). Variants of *THBS1* were reported to be related to autism**^35^** and schizophrenia**^36^**. The protein product of *THBS1* is involved in astrocyte induced synaptogenesis**^37^**, and regulates chain migration of interneuron precursors migrating in the postnatal radial migration stream to the olfactory bulb^38^. Moreover, *THBS1* is an activator of TGFβ signaling, and an inhibitor of pro-angiogenic nitric oxide signaling which plays a role in several cancers and immune-inflammatory conditions.

Variants at Chr14q23.1 were associated with cortical surface area and volume of all regions in the occipital lobe, as well as with thickness, surface area and volume of the middle temporal cortex, banks of the superior temporal sulcus, fusiform, supramarginal and precuneus regions, areas associated with discrimination and recognition of language or visual form. These variants are either intergenic between *KIAA0586*, the product of which is a conserved centrosomal protein essential for ciliogenesis, sonic hedgehog signaling and intracellular organization, and *DACT1*, the product of which is a target for *SIRT1* and acts on the wnt/β-catenin pathway. *KIAA0586* has been associated with Joubert syndrome, another condition associated with abnormal cerebellar development. Other variants are intergenic between *DACT1* and *DAAM1* or intronic in *DAAM1*. *DAAM1* has been associated with occipital lobe volume in a previous GWAS.

Locus 6q22.32 contains various SNPs associated with cortical surface area and volume globally, and also within some frontal, temporal and occipital regions. The SNPs are intergenic between *RSPO3* and *CENPW*. *RSPO3* and *CENPW* have been previously associated with intracranial^39,40^ and occipital lobe volumes. *RSPO3* is an activator of the canonical Wnt signaling pathway and a regulator of angiogenesis.

Chr17q21.31 variants were associated with global cortical surface area and volume and with regions in temporal lobe. These variants are intronic in the genes *PLEKHM1, CRHR1, NSF* and *WNT3*. In previous GWAS analyses, these genes have been associated with general cognitive function^41^ and neuroticism^42,43^. *CRHR1, NSF* and *WNT3* were additionally associated with Parkinson’s disease^44-48^ and intracranial volume^39,40,49^. The *NSF* gene also plays a role in Neuronal Intranuclear Inclusion Disease^50^ and *CRHR1* is involved in anxiety and depressive disorders^51^. This chromosomal region also contains the *MAPT* gene, which plays a role in Alzheimer’s disease, Parkinson’s disease, and frontotemporal dementia^52,53^. The protein product of the gene *ZIC4* is a C2H2 zinc finger transcription factor that has an intraneuronal, non-synaptic expression and auto-antibodies to this protein have been associated with subacute sensory neuronopathy, limbic encephalitis and seizures in patients with breast, small cell lung or ovarian cancers. *ZIC4* null mice have abnormal development of the visual pathway^54^ and heterozygous deletion of the gene has also been associated with a congenital cerebellar (Dandy-Walker) malformation^55^, thus implicating it widely in brain development as well as in neurodegeneration. *C2H2ZF* transcription factors are the most widely expressed transcription factors in eukaryotes and show associations with responses to abiotic (environmental) stressors. Another transcription factor, *FOXC1*, also associated with Dandy-Walker syndrome has been previously shown to be associated with risk of all types of ischemic stroke and with stroke severity. Thus, *ZIC4* might be a biological target worth pursuing to ameliorate neurodegenerative disorders.

We found an exonic SNP within the gene *ENPP2* (Autotaxin) at 8q24.12 to be associated with insular cortical volume. This gene is differentially expressed in the frontal cortex of Alzheimer patients^56^ and in mouse models of Alzheimer disease such as the senescence-accelerated mouse prone 8 strain (SAMP8) mouse. Autotaxin is a dual-function ectoenzyme, which is the primary source of the signaling lipid, lysophosphatidic acid. Besides Alzheimer disease, changes in autotaxin/lysophosphatidic acid signaling have also been shown in diverse brain related conditions such as intractable pain, pruritus, glioblastoma, multiple sclerosis and schizophrenia. In the SAMP8 mouse, improvements in cognition noted after administration of LW-AFC, a putative Alzheimer remedy derived from the traditional Chinese medicinal prescription ‘Liuwei Dihuang’ decoction, are correlated with restored expression of four genes in the hippocampus, one of which is *ENPP2*.

Among the other genetic regions identified, many have been linked to neurological and psychiatric disorders, cognitive functioning, cortical development and cerebral structure (detailed listing in Table S36).

Heritability estimates are, as expected, generally higher in the family-based Austrian Stroke Prevention-Family study (ASPS-Fam) than in the Rotterdam Study (RS) for CTh (average h^2^_ASPS-Fam_=0.52; h^2^_RS_=0.26), CSA (0.62 and 0.30) and CV (0.57 and 0.23). This discrepancy is explained by the different heritability estimation methods: pedigree-based heritability in ASPS-Fam versus heritability based on common SNPs that are in LD with causal variants^57^ in RS. Average heritability over regions is also higher for surface area and volume, than for thickness. The observed greater heritability of CSA compared to CTh is consistent with the previously articulated hypothesis, albeit based on much smaller numbers, that CSA is developmentally determined to a greater extent with smaller subsequent decline after young adulthood, whereas CTh changes over the lifespan as aging, neurodegeneration and vascular injuries accrue^1,3^. It is also interesting that brain regions more susceptible to early amyloid deposition (e.g. superior temporal cortex, precuneus) have a higher heritability. We found no or weak genetic correlation between CTh and CSA, globally and regionally, and no common lead SNPs, which indicates that these two morphological measures are genetically independent, a finding consistent with prior reports^25,26^. In contrast, we found strong genetic correlation between CSA and CV and identified common lead SNPs for CSA and CV globally, and in 12 cortical regions. Similar findings have been reported in a previous publication^26^. The genetic correlation between CTh and CV ranged between 0.09 and 0.77, implying a common genetic background in some regions (such as the primary sensory postcentral and lingual cortices), but not in others. For CTh, we observed genetic correlations between multiple regions within each of the lobes, whereas for CSA and CV we found genetic correlations mainly between different regions of the occipital lobe. Chen et al^58^ have also reported strong genetic correlation for CSA within the occipital lobe. There were also a few genetic correlations observed for regions from different lobes, suggesting similarities in cortical development transcended traditional lobar boundaries.

A limitation of our study is the heterogeneity of the MR phenotypes between cohorts due to different scanners, field strengths, MR protocols and MRI analysis software. Therefore, association results were combined using a sample-size weighted meta-analysis which does not provide overall effect estimates. Moreover, our sample comprises of mainly European ancestry, limiting the generalizability to other ethnicities. Strengths of our study are the population-based design, the large age range of our sample (12 – 90 years), use of three cortical measures as phenotypes of cortical morphometry, and the replication of our CTh and CSA findings in a large and independent cohort. In conclusion, we identified patterns of heritability and genetic associations with various global and regional cortical measures, as well as overlap of MRI cortical measures with genetic traits and diseases that provide new insights into cortical development, morphology and possible mechanisms of disease susceptibility.

## Methods

### Study Population

The sample of this study consist of up to 22,824 participants from 20 population-based cohort studies collaborating in the Cohorts of Heart and Aging Research in Genomic Epidemiology (CHARGE) consortium^59^ and the UK Biobank (UKBB)^60^. All the individuals were stroke- and dementia free, aged between 12 and 90 years, and of European ancestry, except for ARIC AA with African ancestry. Table S1 provides population characteristics of each cohort and Supplementary Section 1 provides a short description of each study. Each study secured approval from institutional review boards or equivalent organizations, and all participants provided written informed consent. Our results were replicated using summary GWAS findings of 22,635 individuals from the Enhancing Neuroimaging Genetics through Meta-analysis (ENIGMA) consortium^31^.

### Genotyping and Imputation

Genotyping was conducted using various commercially available genotyping arrays across the study cohorts. Prior to imputation, extensive quality control was performed in each cohort. Genotype data were imputed to the 1000 Genomes reference panel^61^ (mainly phase 1, version 3) using validated software. Details on genotyping, quality control and imputation can be found in Table S2.

### Phenotype Definition

This study investigated CTh, CSA and CV globally in the whole cortex and in 34 cortical regions. Global and regional CTh was defined as the mean thickness of the left and the right hemisphere in millimeter (mm). Global CSA was defined as the total surface area of the left and the right hemisphere in mm^2^, while regional CSA was defined as the mean surface area of the left and the right hemisphere in mm^2^. Global and regional CV was defined as the mean volume of the left and the right hemisphere in mm^3^. The 34 cortical regions are listed in Table S3. High resolution brain magnetic resonance imaging (MRI) data was obtained in each cohort using a range of MRI scanners, field strengths and protocols. CTh, CSA and CV were generated using the Freesurfer software package^62,63^ in all cohorts except for FHSucd, where an in-house segmentation method was used. MRI protocols of each cohort can be found in Table S4 and descriptive statistics of CTh, CSA and CV can be found in Tables S5, S6 and S7.

### Genome-wide associations, meta-analysis, replication and annotation

Based on a predefined analysis plan, each study fitted linear regression models to determine the association between global and regional CTh, CSA and CV and allele dosages of single nucleotide polymorphisms (SNPs). Additive genetic effects were assumed and the models were adjusted for sex, age, age^2^, and if needed for study site and for principal components to correct for population stratification. Cohorts including related individuals calculated linear mixed models to account for family structure. Details on association software and covariates for each cohort are shown in Table S2. Models investigating regional CTh, CSA and CV were additionally adjusted for global CTh, global CSA and global CV, respectively. Quality control of the summary statistics shared by each cohort was performed using EasyQC^64^. Genetic variants with a minor allele frequency (MAF) < 0.05, low imputation quality (R^2^<0.4), and which were available in less than 10,000 individuals were removed from the analyses. Details on quality control are provided in Supplementary Section 2.

We then used METAL^65^ to perform meta-analyses using the z-scores method, based on p-values, sample size and direction of effect, with genomic control correction. We performed 10,000 permutation tests based on cortical measurements from Rotterdam Study to estimate the number of independent tests. Based on the permutation test results, the genome-wide significance threshold was set a priori at 1.09×10-9 (= 5×10^-8^ /46). We used the clumping function in PLINK^66^ (linkage disequilibrium (LD) threshold: 0.2, distance: 300kb) to identify the most significant SNP in each LD block.

For replication of our genome-wide significant CTh and CSA associations, we used GWAS meta-analysis results from the ENIGMA consortium^31^ for all SNPs that were associated at a p-value < 5×10^-8^ and performed a pooled meta-analysis. The p-value threshold for replication was set to 3.1×10^-4^ (nominal significance threshold (0.05) divided by total number of lead SNPs (160)). CV was not available in the ENIGMA results. The NHGRI-EBI Catalog of published GWAS^67^ was searched for previous SNP-trait associations at a p-value of 5×10^-8^ of lead SNPs.

Regional association plots were generated with LocusZoom^68^, and the chromosomal ideogram with PHENOGRAM (http://visualization.ritchielab.org/phenograms/plot). Annotation of genome-wide significant variants was performed using the ANNOVAR software package^69^ and the FUMA web application^70^. FUMA eQTL mapping uses information from three data repositories (GTEx, Blood eQTL browser, and BIOS QTL browser) and maps SNPs to genes based on a significant eQTL association. We used a false discovery rate threshold (FDR) of 0.05 divided by number of tests (46) to define significant eQTL associations. Gene-based analyses, to combine the effects of SNPs assigned to a gene, and gene set analyses, to find out if genes assigned to significant SNPs were involved in biological pathways, were performed using MAGMA^71^ as implemented in FUMA. The significance threshold was set to 5.87×10^-8^ for gene-based analyses (FDR threshold (0.05) divided by number of genes (18,522) and number of independent tests (46)) and to 1.02×10^-7^ for the gene-set analyses (FDR threshold (0.05) divided by the number of gene sets (10,651) and by the number of independent tests (46)). Additionally, FUMA was used to investigate a significant chromatin interaction between a genomic region in a risk locus and promoter regions of genes (250 bp upstream and 500 bp downstream of a TSS). We used an FDR of 1×10^-6^ to define significant interactions.

We investigated cis (<1Mb) and trans (>1 MB or on a different chromosome) expression quantitative trait loci (eQTL) for genome-wide significant SNPs in 724 post-mortem brains from the Religious Order Study and the Rush Memory and Aging Project (ROSMAP)^72,73^ stored in the AMP-AD database. The samples were collected from the gray matter of the dorsolateral prefrontal cortex. The significance threshold was set to 0.001 (FDR threshold (0.05) divided by the number of independent tests (46)).

For additional pathway analyses of genes that were mapped to independent lead SNPs by FUMA, we searched the InnateDB database^74^. The STRING database^75^ was used for visualizing protein-protein interactions. Only those protein the subnetworks with five or more nodes are shown.

### Heritability

Additive genetic heritability (h^2^) of CTh, CSA and CV was estimated in two studies: the Austrian Stroke Prevention Family Study (ASPS-Fam; n=365) and the Rotterdam Study (RS, n=4472). In the population based family study ASPS-Fam, the ratio of the genotypic variance to the phenotypic variance was calculated using variance components models in SOLAR^76^. In case of non-normalty, phenotype data were inverse-normal transformed. In RS, SNP-based heritability was computed with GCTA^77,78^. These heritability analyses were adjusted for age and sex.

Heritability and partitioned heritability based on GWAS summary statistics was calculated from GWAS summary statistics using LD score regression (LDSC) implemented in the ldsc tool (https://github.com/bulik/ldsc). Partitioned heritability analysis splits genome-wide SNP heritability into 53 functional annotation classes (e.g. coding, 3’ UTR, promoter, transcription factor binding sites, conserved regions etc.) and additionally to 10 cell-type specific classes (e.g. central nervous system, cardiovascular, liver, skeletal muscle etc.) as defined by Finucane et al.^79^ to estimate their contributions to heritability. The significance threshold was set to 2.05×10^-5^ (0.05/number of functional annotation classes (53) / number of independent tests (46)) for heritability partitioned on functional annotation classes and 2.05<10^-6^ (0.05/number of functional annotation classes (53) / number of cell types (10) / number of independent tests (46)) for heritability partitioned on annotation classes and cell types.

### Genetic correlation

LDSR genetic correlation^80^ between CTh, CSA and CV was estimated globally and within each cortical region. The significance threshold was set to 7.35×10-4 (nominal threshold (0.05) divided by number of regions (34) and by number of correlations (CSA and CV, CSA and CTh). Genetic correlation was also estimated between all 34 cortical regions for CTh, CSA and CV, with the significance threshold set to 8.91×10-5 (nominal threshold (0.05) divided by number of regions (34) times the number of regions −1 (33) divided by 2 (half of the matrix). Additionally, the amount of genetic correlation was quantified between CTh, CSA and CV and physical traits (height, BMI), neurological and psychiatric diseases (e.g. Alzheimer’s disease, Parkinson’s disease), cognitive traits and MRI volumes (p-value threshold (0.05/46/number of GWAS traits). As recommended by the ldst tool developers, only HapMap3 variants were included in these analyses, as these tend to be well-imputed across cohorts.

## CHARGE

Infrastructure for the CHARGE Consortium is supported in part by the National Heart, Lung, and Blood Institute grant HL105756 and for the neuroCHARGE phenotype working group through the National Institute on Aging grant AG033193.

### Atherosclerosis Risk in Communities Study (ARIC)

The Atherosclerosis Risk in Communities (ARIC) study is carried out as a collaborative study supported by the National Heart, Lung, and Blood Institute (NHLBI) contracts (HHSN268201100005C, HHSN268201100006C, HHSN268201100007C, HHSN268201100008C, HHSN268201100009C, HHSN268201100010C, HHSN268201100011C, and HHSN268201100012C). The authors thank the staff and participants of the ARIC study for their important contributions. Funding support for “Building on GWAS for NHLBI-diseases: the U.S. CHARGE consortium” was provided by the NIH through the American Recovery and Reinvestment Act of 2009 (ARRA) (5RC2HL102419). This project was also funded from R01-NS087541.

### Austrian Stroke Prevention Family (ASPS) / Austrian Stroke Prevention Family Study (ASPS-Fam)

The authors thank the staff and the participants for their valuable contributions. We thank Birgit Reinhart for her long-term administrative commitment, Elfi Hofer for the technical assistance at creating the DNA bank, Ing. Johann Semmler and Anita Harb for DNA sequencing and DNA analyses by TaqMan assays and Irmgard Poelzl for supervising the quality management processes after ISO9001 at the biobanking and DNA analyses. The Medical University of Graz and the Steiermärkische Krankenanstaltengesellschaft support the databank of the ASPS/ASPS-Fam. The research reported in this article was funded by the Austrian Science Fund (FWF) grant numbers PI904, P20545-P05 and P13180 and supported by the Austrian National Bank Anniversary Fund, P15435 and the Austrian Ministry of Science under the aegis of the EU Joint Programme-Neurodegenerative Disease Research (JPND)-www.jpnd.eu.

### Cardiovascular Health Study (CHS)

Cardiovascular Health Study: This CHS research was supported by NHLBI contracts HHSN268201200036C, HHSN268200800007C, HHSN268201800001C, N01HC55222, N01HC85079, N01HC85080, N01HC85081, N01HC85082, N01HC85083, N01HC85086, N01HC15103, HHSN268200960009C; and NHLBI grants U01HL080295, R01HL087652, R01HL105756, R01HL103612, R01HL120393, and U01HL130114 with additional contribution from the National Institute of Neurological Disorders and Stroke (NINDS). Additional support was provided through R01AG023629, and R01AG033193 from the National Institute on Aging (NIA). A full list of principal CHS investigators and institutions can be found at CHS-NHLBI.org.

The provision of genotyping data was supported in part by the National Center for Advancing Translational Sciences, CTSI grant UL1TR001881, and the National Institute of Diabetes and Digestive and Kidney Disease Diabetes Research Center (DRC) grant DK063491 to the Southern California Diabetes Endocrinology Research Center. The content is solely the responsibility of the authors and does not necessarily represent the official views of the National Institutes of Health.

### Erasmus Rucphen Family Study (ERF)

Erasmus Rucphen Family (ERF) was supported by the Consortium for Systems Biology (NCSB), both within the framework of the Netherlands Genomics Initiative (NGI)/Netherlands Organisation for Scientific Research (NWO). ERF study as a part of EUROSPAN (European Special Populations Research Network) was supported by European Commission FP6 STRP grant number 018947 (LSHG-CT-2006-01947) and also received funding from the European Community’s Seventh Framework Programme (FP7/2007-2013)/grant agreement HEALTH-F4-2007-201413 by the European Commission under the programme “Quality of Life and Management of the Living Resources” of 5th Framework Programme (No. QLG2-CT-2002-01254) as well as FP7 project EUROHEADPAIN (nr 602633). High-throughput analysis of the ERF data was supported by joint grant from Netherlands Organisation for Scientific Research and the Russian Foundation for Basic Research (NWO-RFBR 047.017.043). High throughput metabolomics measurements of the ERF study has been supported by BBMRI-NL (Biobanking and Biomolecular Resources Research Infrastructure Netherlands).

### Framingham Heart Study (FHS)

This work was supported by the National Heart, Lung and Blood Institute’s Framingham Heart Study (Contract No. N01-HC-25195 and No. HHSN268201500001I) and its contract with Affymetrix, Inc. for genotyping services (Contract No. N02-HL-6-4278). A portion of this research utilized the Linux Cluster for Genetic Analysis (LinGA-II) funded by the Robert Dawson Evans Endowment of the Department of Medicine at Boston University School of Medicine and Boston Medical Center. This study was also supported by grants from the National Institute of Aging (R01s AG033040, AG033193, AG054076, AG049607, AG008122, AG016495; and U01-AG049505) and the National Institute of Neurological Disorders and Stroke (R01-NS017950). We would like to thank the dedication of the Framingham Study participants, as well as the Framingham Study team, especially investigators and staff from the Neurology group, for their contributions to data collection. Dr. DeCarli is supported by the Alzheimer’s Disease Center (P30 AG 010129). The views expressed in this manuscript are those of the authors and do not necessarily represent the views of the National Heart, Lung, and Blood Institute; the National Institutes of Health; or the U.S. Department of Health and Human Services.

### Lothian Birth Cohort 1936 (LBC1936)

This project is funded by the Age UK’s Disconnected Mind programme (http://www.disconnectedmind.ed.ac.uk) and also by Research Into Ageing (Refs. 251 and 285). The whole genome association part of the study was funded by the Biotechnology and Biological Sciences Research Council (BBSRC; Ref. BB/F019394/1). Analysis of the brain images was funded by the Medical Research Council Grants G1001401 and 8200 and MR/M01311/1. The imaging was performed at the Brain Research Imaging Centre, The University of Edinburgh (http://www.bric.ed.ac.uk), a centre in the SINAPSE Collaboration (http://www.sinapse.ac.uk). The work was undertaken by The University of Edinburgh Centre for Cognitive Ageing and Cognitive Epidemiology (http://www.ccace.ed.ac.uk), part of the cross council Lifelong Health and Wellbeing Initiative (Ref. G0700704/84698). Funding from the BBSRC, Medical Research Council (MR/K026992/1) and Scottish Funding Council through the SINAPSE Collaboration is gratefully acknowledged. We thank the LBC1936 participants and research team members. We also thank the nurses and staff at the Wellcome Trust Clinical Research Facility (http://www.wtcrf.ed.ac.uk), where subjects were tested and the genotyping was performed.

### LIFE-Adult

LIFE-Adult is funded by the Leipzig Research Center for Civilization Diseases (LIFE). LIFE is an organizational unit affiliated to the Medical Faculty of the University of Leipzig. LIFE is funded by means of the European Union, by the European Regional Development Fund (ERDF) and by funds of the Free State of Saxony within the framework of the excellence initiative. This work was also funded by the Deutsche Forschungsgemeinschaft (Grant Number: CRC 1052 “Obesity mechanisms” project A1 to AV) and by the Max Planck Society.

### Sydney Memory and Ageing Study (MAS)

MAS is funded by the Australian National Health and Medical Research Council (NHMRC)/Australian Research Council Strategic Award (Grant 401162), NHMRC Project grant 1405325.

We would like to gratefully acknowledge and thank the Sydney MAS participants and supporters and the Sydney MAS Research Team.

### Older Australian Twin Study (OATS)

OATS is funded by the Australian National Health and Medical Research Council (NHMRC)/Australian Research Council Strategic Award (Grant 401162), NHMRC Program Grants (350833, 568969, 109308)

We would like thank and gratefully acknowledge the OATS participants, their supporters and the OATS Research Team.

### Rotterdam Study (RSI, RSII, RSIII)

The Rotterdam Study is funded by Erasmus Medical Center and Erasmus University, Rotterdam, Netherlands Organization for the Health Research and Development (ZonMw), the Research Institute for Diseases in the Elderly (RIDE), the Ministry of Education, Culture and Science, the Ministry for Health, Welfare and Sports, the European Commission (DG XII), and the Municipality of Rotterdam. The authors are grateful to the study participants, the staff from the Rotterdam Study and the participating general practitioners and pharmacists. The generation and management of GWAS genotype data for the Rotterdam Study (RS I, RS II, RS III) were executed by the Human Genotyping Facility of the Genetic Laboratory of the Department of Internal Medicine, Erasmus MC, Rotterdam, The Netherlands. The GWAS datasets are supported by the Netherlands Organisation of Scientific Research NWO Investments (nr. 175.010.2005.011, 911-03-012), the Genetic Laboratory of the Department of Internal Medicine, Erasmus MC, the Research Institute for Diseases in the Elderly (014-93-015; RIDE2), the Netherlands Genomics Initiative (NGI)/Netherlands Organisation for Scientific Research (NWO) Netherlands Consortium for Healthy Aging (NCHA), project nr. 050-060-810. We thank Pascal Arp, Mila Jhamai, Marijn Verkerk, Lizbeth Herrera and Marjolein Peters, and Carolina Medina-Gomez, for their help in creating the GWAS database, and Karol Estrada, Yurii Aulchenko, and Carolina Medina-Gomez, for the creation and analysis of imputed data. This work has been performed as part of the CoSTREAM project (www.costream.eu) and ORACLE project, and has received funding from the European Union’s Horizon 2020 research and innovation programme under grant agreement No 667375 and No 678543.

### Study of Health in Pomerania (SHIP) / Study of Health in Pomerania Trend (SHIP-Trend)

SHIP is part of the Community Medicine Research net of the University of Greifswald, Germany, which is funded by the Federal Ministry of Education and Research (grants no. 01ZZ9603, 01ZZ0103, and 01ZZ0403), the Ministry of Cultural Affairs as well as the Social Ministry of the Federal State of Mecklenburg-West Pomerania, and the network ‘Greifswald Approach to Individualized Medicine (GANI_MED)’ funded by the Federal Ministry of Education and Research (grant 03IS2061A). Genome-wide data have been supported by the Federal Ministry of Education and Research (grant no. 03ZIK012) and a joint grant from Siemens Healthineers, Erlangen, Germany and the Federal State of Mecklenburg- West Pomerania. Whole-body MR imaging was supported by a joint grant from Siemens Healthineers, Erlangen, Germany and the Federal State of Mecklenburg West Pomerania. The University of Greifswald is a member of the Caché Campus program of the InterSystems GmbH.

### Saguenay Youth Study (SYS)

The Saguenay Youth Study has been funded by the Canadian Institutes of Health Research (TP, ZP), Heart and Stroke Foundation of Canada (ZP), and the Canadian Foundation for Innovation (ZP). We thank all families who took part in the Saguenay Youth Study. SYS is supported by the Canadian Institutes of Health Research: NET54015, NRF86678, TMH109788.

### Three-City Dijon (3C-Dijon)

The Three City (3C) Study is conducted under a partnership agreement among the Institut National de la Santé et de la Recherche Médicale (INSERM), the University of Bordeaux, and Sanofi-Aventis. The Fondation pour la Recherche Médicale funded the preparation and initiation of the study. The 3C Study is also supported by the Caisse Nationale Maladie des Travailleurs Salariés, Direction Générale de la Santé, Mutuelle Générale de l’Education Nationale (MGEN), Institut de la Longévité, Conseils Régionaux of Aquitaine and Bourgogne, Fondation de France, and Ministry of Research–INSERM Programme “Cohortes et collections de données biologiques.” Christophe Tzourio and Stéphanie Debette have received investigator-initiated research funding from the French National Research Agency (ANR) and from the Fondation Leducq. Stéphanie Debette is supported by a starting grant from the European Research Council (SEGWAY) and a grant from the Joint Programme of Neurodegenerative Disease research (BRIDGET), from the European Union’s Horizon 2020 research and innovation programme under grant agreements No 643417 & No 640643, and by the Initiative of Excellence of Bordeaux University. We thank Dr. Anne Boland (CNG) for her technical help in preparing the DNA samples for analyses. This work was supported by the National Foundation for Alzheimer’s disease and related disorders, the Institut Pasteur de Lille, the labex DISTALZ and the Centre National de Génotypage.

### United Kingdom Biobank (UKBB)

This research has been conducted using the UK Biobank Resource under Application Number “23509”.

### Vietnam Era Twin Study of Aging (VETSA)

This work was supported by US National Institutes of Health grants AG018386, AG022381, AG022982, AG050595, AG018384, AG046413, AG047903, DA025109, DA023549, DA18673, HD050735, and U 54EB020403, and the VA San Diego Center of Excellence for Stress and Mental Health Healthcare System. The content is the responsibility of the authors and does not necessarily represent official views of the NIA, NIH, or VA. The Cooperative Studies Program of the U.S. Department of Veterans Affairs provided financial support for development and maintenance of the Vietnam Era Twin Registry. We would also like to acknowledge the continued cooperation and participation of the members of the VET Registry and their families.

## ENIGMA

The study was supported in part by grant U54 EB020403 from the NIH Big Data to Knowledge (BD2K) Initiative, a cross-NIH partnership. Additional support was provided by R01MH116147, P41 EB015922, RF1AG051710, RF1 AG041915 (to P.T.), by P01 AG026572, R01 AG059874 and by R01 MH117601 (to N.J. and L.S.). S.E.M. was funded by an NHMRC Senior Research Fellowship (APP1103623). L.C.-C. was supported by a QIMR Berghofer Fellowship.

### Author Contributions

#### Drafting of the Manuscript

Edith Hofer, Gennady V. Roshchupkin, Reinhold Schmidt, Sudha Seshadri

#### Genotype Data Acquisition

Helena Schmidt, Najaf Amin, Peter R Schofield, Margaret J Wright, Carol E. Franz, Margaret J Wright

#### Phenotype Data Acquisition

Reinhold Schmidt, Sophie Maingault., Bernard Mazoyer, Jiyang Jiang, Julian Trollor, G. Bruce Pike, Carol E. Franz

#### Genetic Data Analysis and Bioinformatics Analysis

Edith Hofer, Yasaman Saba, Rui Xia, Josh C. Bis, Shahzad Ahmad, Shuo Li, Sven J van der Lee, Qiong Yang, Claudia Satizabal, Honghuang Lin, Jayandra J Himali, Habil Zare, Michelle Luciano, Markus Scholz, Nicola J. Armstrong, Gennady V. Roshchupkin, Hieab H. H. Adams, Maria J Knol, Meike W Vernooij, Alexander Teumer, Katharina Wittfeld, Manon Bernard, Aniket Mishra, Nathan A. Gillespie, Mark Logue, Peter R Schofield

#### Imaging Data Analysis

Gennady V. Roshchupkin, Hieab H. H. Adams, Lukas Pirpamer, Stephan Seiler, Pauline Maillard, Charles DeCarli, Sherif Karama, Lindsay Lewis, Mark Bastin, Wei Wen, Frauke Beyer, A. Veronica Witte, Mat Harris, Jiyang Jiang

#### Cohort PIs

Perminder S. Sachdev, William S. Kremen, Joanna A. Wardlaw, Arno Villringer, Cornelia M. van Duijn, Hans Jörgen Grabe, William T. Longstreth Jr, Myriam Fornage, Tomas Paus, Stephanie Debette, M. Arfan Ikram, Helena Schmidt, Reinhold Schmidt, Sudha Seshadri, Jerome I Rotter, Bruce M Psaty, Ian J Deary, Markus Loeffler, Albert Hofman, André G Uitterlinden, Wiro J Niessen, Zdenka Pausova, Matthew S. Panizzon, Thomas H Mosley, Julian Trollor, Carol E. Franz

#### ENIGMA Study Design

Katrina L. Grasby, Neda Jahanshad, Jodie N. Painter, Lucía Colodro-Conde, Janita Bralten, Derrek P. Hibar, Penelope A. Lind, Fabrizio Pizzagalli,Jason L. Stein, Paul M. Thompson, Sarah E. Medland

