## Supplementary Information for "Genetic Determinants of Cortical Structure (Thickness, Surface Area and Volumes) among Disease Free Adults in the CHARGE Consortium"


1. **Cohorts**
2. **Quality Control**
3. **Supplementary Figures**

   **Supplementary Figure 1.** Regional association plots for global cortical surface area.

   **Supplementary Figure 2.** Regional association plots for global cortical volume.

   **Supplementary Figure 3.** Regional association plots for postcentral cortical thickness, surface area and volume.

   **Supplementary Figure 4.** QQ plots of global cortical thickness, surface area and volume meta-analyses.

   **Supplementary Figure 5.** QQ plots of regional cortical thickness meta-analyses.

   **Supplementary Figure 6.** QQ plots of regional cortical surface area meta-analyses.

   **Supplementary Figure 7.** QQ plots of regional cortical volume meta-analyses.

   **Supplementary Figure 8.** Pathway analysis of the 44 genes that mapped to independent lead SNPs of cortical thickness.

   **Supplementary Figure 9.** Pathway analysis of the 105 genes that mapped to independent lead SNPs of cortical surface area.

   **Supplementary Figure 10.** Pathway analysis of the 82 genes that mapped to independent lead SNPs of cortical volume.

   **Supplementary Figure 11.** Regional heritability estimates based on common SNPs.

   **Supplementary Figure 12.** Genetic correlation between cortical thickness, surface area and volume within cortical regions.

   **Supplementary Figure 13.** Genetic correlation between regional cortical thickness.

   **Supplementary Figure 14.** Genetic correlation between regional cortical surface area.

   **Supplementary Figure 15.** Genetic correlation between regional cortical volume.

   **Supplementary Figure 16.** Genetic correlation between cortical thickness and other GWAS phenotypes.

   **Supplementary Figure 17.** Genetic correlation between cortical surface area and other GWAS phenotypes.

   **Supplementary Figure 18.** Genetic correlation between cortical volume and other GWAS phenotypes.
4. **References**

### 1. Cohorts

**Atherosclerosis Risk in Communities  Study (ARIC)**

The ARIC study is a population-based cohort study of atherosclerosis and clinical atherosclerotic diseases[1]. At its inception (1987-1989), 15,792 men and women, including 11,478 white and 4,266 black participants were recruited from four U.S. communities: Suburban Minneapolis, Minnesota; Washington County, Maryland; Forsyth County, North Carolina; and Jackson, Mississippi. In the first 3 communities, the sample reflects the demographic composition of the community. In Jackson, only black residents were enrolled. Participants were between age 45 and 64 years at their baseline examination in 1987-1989 when blood was drawn for DNA extraction and participants consented to genetic testing. Vascular risk factors and outcomes, including transient ischemic attack, stroke and dementia, were determined in a standard fashion. During the first 2 years (1993-1994) of the third ARIC examination (V3), participants aged 55 and older from the Forsyth County and Jackson sites were invited to undergo cranial MRI. This subgroup of individuals with MRI scanning represents a random sample of the full cohort because examination dates were allocated at baseline through randomly selected induction cycles.

**Austrian Stroke Prevention Study (ASPS)**

The ASPS study is a single center prospective follow-up study on the effects of vascular risk factors on brain structure and function in the normal elderly population of the city of Graz, Austria. The procedure of recruitment and diagnostic work-up of study participants has been described previously[2,3]. A total of 2007 participants were randomly selected from the official community register stratified by gender and 5 year age groups. Individuals were excluded from the study if they had a history of neuropsychiatric disease, including previous stroke, transient ischemic attacks, and dementia, or an abnormal neurologic examination determined on the basis of a structured clinical interview and a physical and neurologic examination. During 2 study periods between September 1991 and March 1994 and between January 1999 and December 2003 an extended diagnostic work-up including MRI and neuropsychological testing was done in 1076 individuals aged 45 to 85 years randomly selected from the entire cohort: 509 from the first period and 567 from the second. In 1992, blood was drawn from all study participants for DNA extraction. They were all European Caucasians. Genotyping was performed in 996 participants, and the 182 who underwent MRI scanning at baseline were available for these analyses.

**Austrian Stroke Prevention Family Study (ASPS-Fam)**

ASPS-Fam is a prospective single-center community-based study on the cerebral effects of vascular risk factors in the normal aged population of the city of Graz, Austria[4,5]. ASPS-Fam represents an extension of the Austrian Stroke Prevention Study (ASPS), which was established in 1991[2,3]. Between 2006 and 2013, study participants of the ASPS and their first-grade relatives were invited to enter ASPS-Fam.

Inclusion criteria were no history of previous stroke or dementia and a normal neurologic examination. A total of 419 individuals from 176 families were included into the study. The number of members per family ranged from 2 to 6. The entire cohort underwent a thorough diagnostic workup including clinical history, laboratory evaluation, cognitive testing, and an extended vascular risk factor assessment. They were all European Caucasians. Those 297 participants who passed genotyping quality control and underwent MRI scanning were available for these analyses.

**Cardiovascular Health Study (CHS)**

The CHS is a population-based cohort study of risk factors for coronary heart disease and stroke in adults ≥ 65 years conducted across four field centers[6]. The original predominantly European ancestry cohort of 5,201 persons was recruited in 1989-1990 from random samples of the Medicare eligibility lists; subsequently, an additional predominantly African-American cohort of 687 persons was enrolled for a total sample of 5,888. Blood samples were drawn from all participants at their baseline examination and DNA was subsequently extracted from available samples. Genotyping was performed at the General Clinical Research Center's Phenotyping/Genotyping Laboratory at Cedars-Sinai among CHS participants who consented to genetic testing and had DNA. European ancestry participants were excluded from the GWAS study sample due to the presence at study baseline of coronary heart disease, congestive heart failure, peripheral vascular disease, valvular heart disease, stroke or transient ischemic attack or lack of available DNA. Among those with successful GWAS, 567 European ancestry participants had available FreeSurfer measures for this analysis. CHS was approved by institutional review committees at each field center and individuals in the present analysis had available DNA and gave informed consent including consent to use of genetic information for the study of cardiovascular disease.

**Erasmus Rucphen Family  Study (ERF)**

The Erasmus Rucphen Family genetic isolate study (ERF) is a prospective family based study located in Southwest of the Netherlands[7,8]. This young genetic isolate was founded in the mid-eighteenth century and minimal immigration and marriages occurred between surrounding settlements due to social and religious reasons. The ERF study population includes 3465 individuals that are living descendants of 22 couples with at least six children baptized. Informed consent has been obtained from patients where appropriate. The study protocol was approved by the medical ethics board of the Erasmus Medical Center Rotterdam, the Netherlands.

**Framingham Heart Study (FHS)**

The FHS is a three-generation, single-site, community-based, ongoing cohort study that was initiated in 1948 to investigate the risk factors for cardiovascular disease. It now comprises 3 generations of participants: the Original cohort followed since 1948[9]; their Offspring and spouses of the Offspring (Gen 2), followed since 1971[10]; and children from the largest Offspring families enrolled in 2000 (Gen

3)[11]. The Original cohort enrolled 5,209 men and women who comprised two-thirds of the adult population then residing in Framingham, MA. Survivors continue to receive biennial examinations. The Offspring cohort comprises 5,124 persons (including 3,514 biological offspring) who have been examined approximately once every 4 years. The Third-generation includes 4,095 participants with at least one parent in the Offspring Cohort. The first two generations were invited to undergo an initial brain MRI in 1999-2005, and for Gen 3, brain MRI began in 2009. The population of Framingham was virtually entirely white (Europeans of English, Scots, Irish and Italian descent) in 1948 when the Original cohort was recruited. Self-reports of ethnicity across all three generations were 99.7% whites, reflecting the ethnicity of the population of Framingham in 1948. FHS participants had DNA extracted and provided consent for genotyping, and eligible participants underwent genome-wide genotyping.

**Lothian Birth Cohort 1936 (LBC1936)**

The LBC1936 consists of relatively healthy individuals assessed on cognitive and medical measures at age 70 years (n=1,091), and again with brain imaging traits at 73 years of age (n=866). They were born in 1936, most took part in the Scottish Mental Survey of 1947, and almost all lived independently in the Lothian region of Scotland. A full description of participant recruitment and testing can be found elsewhere[12-14]. The study was approved by the Lothian (REC 07/MRE00/58) and Scottish Multicentre (MREC/01/0/56) Research Ethics Committees and all subjects give written informed consent.

**LIFE-Adult**

LIFE-Adult is a population-based study and a part of the large-scale research project LIFE (Leipzig Research Center for Civilization Diseases). 10,000 residents (main age range 40 – 79) from the district of Leipzig (Saxony, Germany) were recruited and extensively phenotyped for a number of disease and environmental parameters (see Loeffler et al.[15] for a description of the assessment programme). All subjects gave written informed consent to participate in the study. The procedures were conducted according to the Declaration of Helsinki and approved by the University of Leipzig's ethics committee (registration-number: 263-2009-14122009).

**Sydney Memory and Ageing Study (MAS)**

The Sydney Memory and Ageing Study recruited participants aged 70-90 years of age randomly from the community in Sydney, Australia (N=1037 at baseline). Exclusion criteria included a diagnosis of dementia, schizophrenia or a progressive malignancy. Comprehensive data was collected, including an extensive cognitive battery, demographics and medical history. Most participants provided a blood sample for genetic and biochemistry analyses. A subset of participants also underwent neuroimaging. Ethics approval for the study was provided by the University of New South Wales and the Illawarra Area Health Service Human Research Ethics Committees. All participants gave written informed consent. More information is provided in Sachdev et al. 2010[16].

**Older Australian Twin Study (OATS)**

Twins aged 65 years and over were recruited from Twins Registry Australia and through a recruitment drive from the three eastern Australian states into the Older Australian Twins Study (N=623 at baseline). Exclusion criteria included insufficient English to finish the assessment. A comprehensive assessment was undertaken including an extensive cognitive battery and blood was collected for genetic and biochemistry analyses. For a subset, neuroimaging was also performed. Ethics approval for the study was provided by Twins Registry Australia, University of New South Wales, University of Melbourne, Queensland Institute of Medical Research and the South Eastern Sydney and Illawarra Area Health Service University of New South Wales and the Illawarra Area Health Service Human Research Ethics Committees. All participants gave written informed consent. More information is provided in Sachdev et al. 2009[17].

**Rotterdam Study (RSI, RSII, RSIII)**

The Rotterdam Study is a prospective, population-based cohort study among individuals living in the well-defined Ommoord district in the city of Rotterdam in The Netherlands[18]. The aim of the study is to determine the occurrence of cardiovascular, neurological, ophthalmic, endocrine, hepatic, respiratory, and psychiatric diseases in elderly people. The cohort was initially defined in 1990 among approximately 7,900 persons, aged 55 years and older, who underwent a home interview and extensive physical examination at the baseline and during follow-up rounds every 3-4 years (RS-I). The cohort was extended in 2000/2001 (RS-II, 3,011 individuals aged 55 years and older) and 2006/2008 (RS-III, 3,932 subjects, aged 45 and older). Written informed consent was obtained from all participants and the Medical Ethics Committee of the Erasmus Medical Center, Rotterdam, approved the study.

**Study of Health in Pomerania (SHIP)**

The Study of Health in Pomerania (SHIP) is a population-based project in West Pomerania, the north-east area of Germany.[19] A sample from the population aged 20 to 79 years was drawn from population registries. First, the three cities of the region (with 17,076 to 65,977 inhabitants) and the 12 towns (with 1,516 to 3,044 inhabitants) were selected, and then 17 out of 97 smaller towns (with less than 1,500 inhabitants), were drawn at random. Second, from each of the selected communities, subjects were drawn at random, proportional to the population size of each community and stratified by age and gender. Only individuals with German citizenship and main residency in the study area were included. Finally, 7,008 subjects were sampled, with 292 persons of each gender in each of the twelve five-year age strata. In order to minimize drop-outs by migration or death, subjects were selected in two waves. The net sample (without migrated or deceased persons) comprised 6,267 eligible subjects. Selected persons received a maximum of three written invitations. In case of non-response, letters were followed by a phone call or by home visits if contact by phone was not possible. The SHIP population finally comprised 4,308 participants (corresponding to a final response of 68.8%).

**Study of Health in Pomerania Trend (SHIP-Trend)**

The Study of Health in Pomerania (SHIP-Trend) is a population-based cohort study in West Pomerania, a region in the northeast of Germany, assessing the prevalence and incidence of common population-relevant diseases and their risk factors.[19] Baseline examinations for SHIP-Trend were carried out between 2008 and 2012, comprising 4,420 participants aged 20 to 81 years. Study design and sampling methods were previously described. The medical ethics committee of the University of Greifswald approved the study protocol, and oral and written informed consents were obtained from each of the study participants.

**Saguenay Youth Study (SYS)**

The Saguenay Youth Study (SYS) is a two-generational study of adolescents and their parents (n=1,029 adolescents and 962 parents) aimed at investigating the etiology, early stages and trans-generational trajectories of common cardio-metabolic and brain diseases. The cohort was recruited from the genetic founder population of the Saguenay Lac St. Jean region of Quebec, Canada. The participants underwent extensive (15-hour) phenotyping, including an hour-long recording of beat-by-beat blood pressure, magnetic resonance imaging of the brain and abdomen, and serum lipidomic profiling with LC-ESI-MS. All participants have been genome-wide genotyped (with ~8M imputed SNPs) and a subset of them (144 adolescents and their 288 parents) has been genome-wide epityped (whole blood DNA, Infinium HumanMethylation450K BeadChip). These assessments are complemented by a detailed evaluation of each participant in a number of domains, including cognition, mental health and substance use, diet, physical activity and sleep, and family environment. The data collection took place during 2003-2012 in adolescents (full) and their parents (partial), and during 2012-2015 in parents (full). The Research Ethics Committee of the Chicoutimi Hospital approved the study protocol. Written informed consent was also obtained from all the participants.

**Three-City Dijon (3C-Dijon)**

The 3C study is conducted in three French cities (Bordeaux, Dijon, and Montpellier), comprising 9,294 participants, designed to estimate the risk of dementia and cognitive impairment attributable to vascular factors.1 Eligibility criteria included living in the city and being registered on the electoral rolls in 1999, 65 years or older, and not institutionalized. The 3C-Dijon study recruited 4,931 individuals. The overall design of the 3C-Dijon study is detailed elsewhere[20-22]. Participants aged less than 80 years and enrolled between June 1999 and September 2000 (n=2,763) were invited to undergo a brain MRI. Although 2,285 subjects agreed to participate (82.7%), because of financial limitations, 1,924 MRI scans were performed. DNA samples of 3C-Dijon participants were genotyped at the Centre National de Génotypage, Evry, France, with Illumina Human610IQuad® BeadChips.

**United Kingdom Biobank (UKBB)**

The UKBB is a large-scale epidemiological study of over 500,000 individuals aged 40-69 years from the

United Kingdom (http://www.ukbiobank.ac.uk). The analyses presented here use data that were accessed via application 1155. Genetic data are available for the majority of these individuals[23] and as of 15 July 2017 13,269 of these participants had participated in a multimodal imaging sub-study [24,25]. The analyses presented here use the cortical measurements of the 8.213 participants released by the UKBB which are derived from the T1 Brain MRI, the extraction of these measures using FreeSurfer software version 6.0. These data were not visually QCed (as the required files were not available for download). However, we removed outliers by setting data points more than 3 standard deviations from the mean to missing. The genetic data used for these analyses uses only those variants imputed using the HRC reference panel. Imputation accuracy and allele frequency were recalculated in the subset of participants with imaging from the raw imputed data using HASE software, quality control filters used in the meta-analyses were applied to the UKBB data prior to analysis. To account for ethnicity, we included only subjects with white British ancestry (base on provided by UK Biobank information). To avoid correct cryptic relationship we excluded all subject with >=3rd degree of genetic relationship.

**Vietnam Era Twin Study of Aging (VETSA)**

Wave 1 of the longitudinal VETSA project comprised middle-aged male twins aged 51-60 years randomly recruited form the population-based Harvard Drug Study (HDS)[1] between 2003 and 2007[2,3]. Eligibility for the HDS was not based on drug use or any clinical or diagnostic characteristic. VETSA participants were concordant for US military service at some time between 1965 and 1975. Nearly 80% reported no combat experience. The sample is 88.3% non-Hispanic white, 5.3% African-American, 3.4% Hispanic, and 3.0% "other" participants. Based on data from the US National Center for Health Statistics, the sample is very similar to American men in their age range with respect to health and lifestyle characteristics[4]. A total of 445 individuals who had genotyping and analyzable MRI data, and were of European ancestry were included in the present analysis. Once generated, the cortical surface model was manually reviewed and edited for technical accuracy. Minimal manual editing was performed by applying standard, objective editing rules. DNA samples were genotyped at deCODE Genetics (Reykjavík, Iceland) with Illumina HumanOmniExpress-24 v1.0A beadchips.

**Enhancing Neuroimaging Genetics through Meta-analysis (ENIGMA)**

The ENIGMA sample includes 22,633 individuals of European ancestry from 44 cohorts. The average age was 39.9 (ranging from 3.4 to 91.4) years. The cohorts were either population-based (23 cohorts) or case-control (21 cohorts), where the cases had a psychiatric or neurological disorder. The cohorts included were: 1000BRAINS, ADNI1, ADNI2GO, BETULA, BIG-Affy, BIG-PsychChip, BONN, BrainScale, CARDIFF, CLING, DNS-V3, DNS-V4, FBIRN, FOR2107, GSP, HUBIN, HUNT, IMAGEN, IMpACT, LBC1936, LIBD, MCIC, MPIP, MPRC, MÜNSTER, NCNG, NESDA, NeuroIMAGE, NTR, OATS, PAFIP, PDNZ, PING, PPMI, QTIM, SHIP, SHIP-Trend, Sydney MAS, SYS, TOP, TOP3T, UiO2016, UiO2017, and UMCU.

#### 2. Quality Control (QC)

**Pre-meta-analyis QC**

We used the protocol implemented in EasyQC[30] to perform quality control before meta-analysis. SNPs were filtered out based on the following QC criteria:

- missing information (alleles, p-value, effect size, standard error of effect-size, frequency of effect allele, sample size for the SNP, imputation quality)
- impossible values (p-value<0 or p-value>1, standard error of effect-size <=0 or standard error of effect-size = infinity, frequency of effect allele <0 or frequency of effect allele > 1, imputation quality<0)
- non A/C/G/T/I/D markers
- duplicated SNPs (the SNP with the lower sample size was removed)
- monomorphic SNPs
- allele mismatches between input and reference (e.g. A/T in input, A/G in reference)
- allele frequency outliers: remove SNPs where |Freq-Freq.ref|>0.2
- number of individuals in the cohort with this SNP <100
- minor allele count<= 6
- imputation quality < 0.4

**Post-meta-analyis QC**

After the meta-analysis we removed SNPs with a minor allele frequency (MAF) < 0.05 and which were available in less than 10000 individuals.

#### 3. Supplementary Figures

**Supplementary Figure 1. Regional association plots for global cortical surface area.**

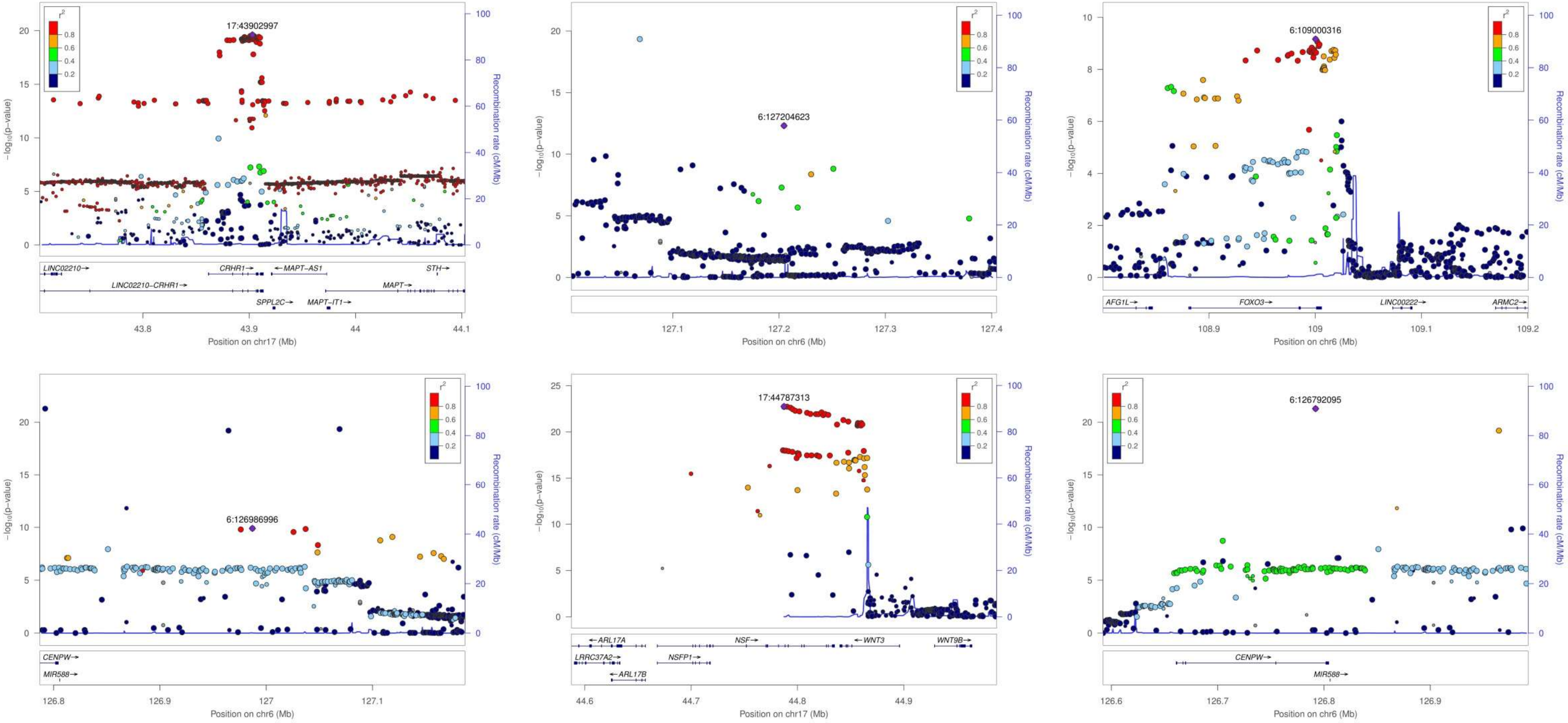

**Supplementary Figure 2. Regional association plots for global cortical volume.**

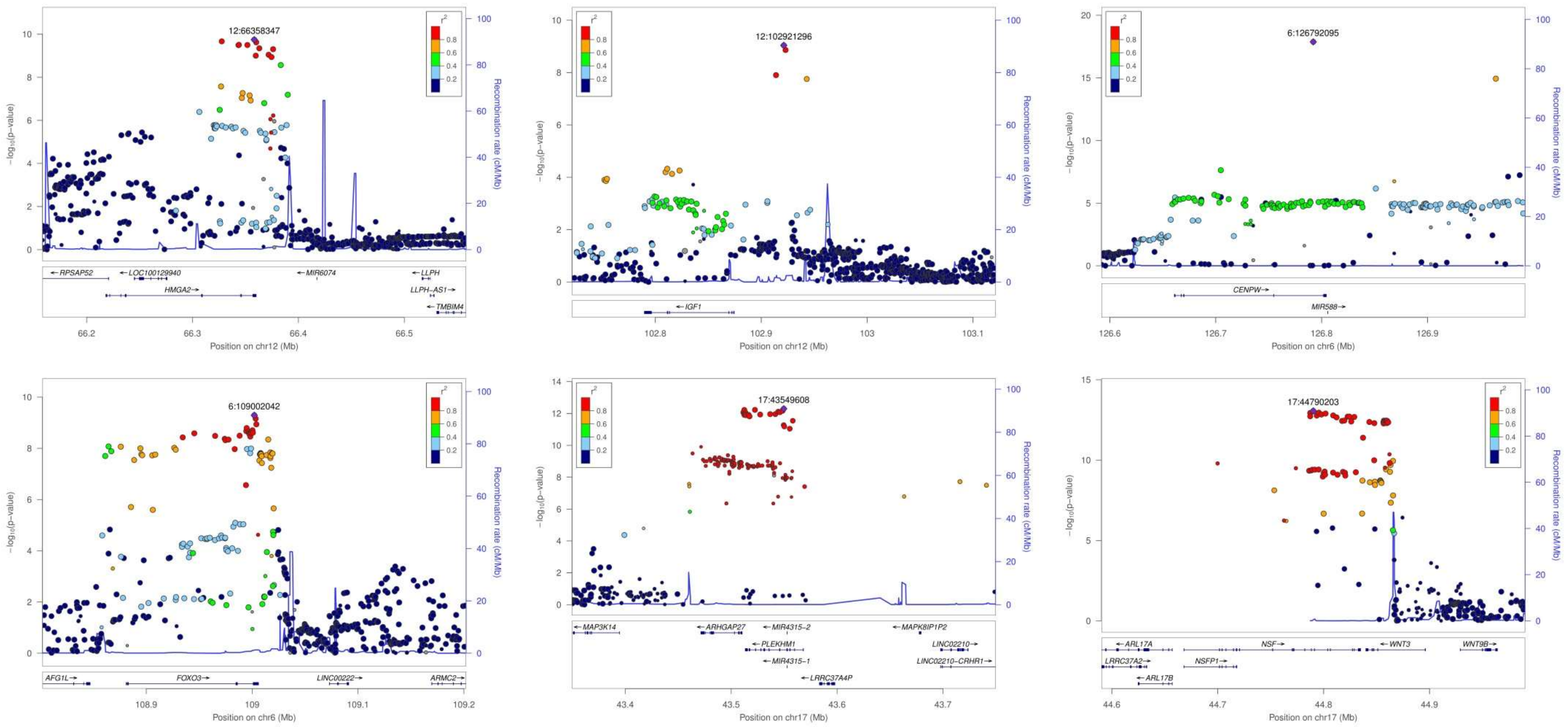

**Supplementary Figure 3. Regional association plots of postcentral cortical thickness (CTh), surface area (CSA) and volume (CV).**

CTh

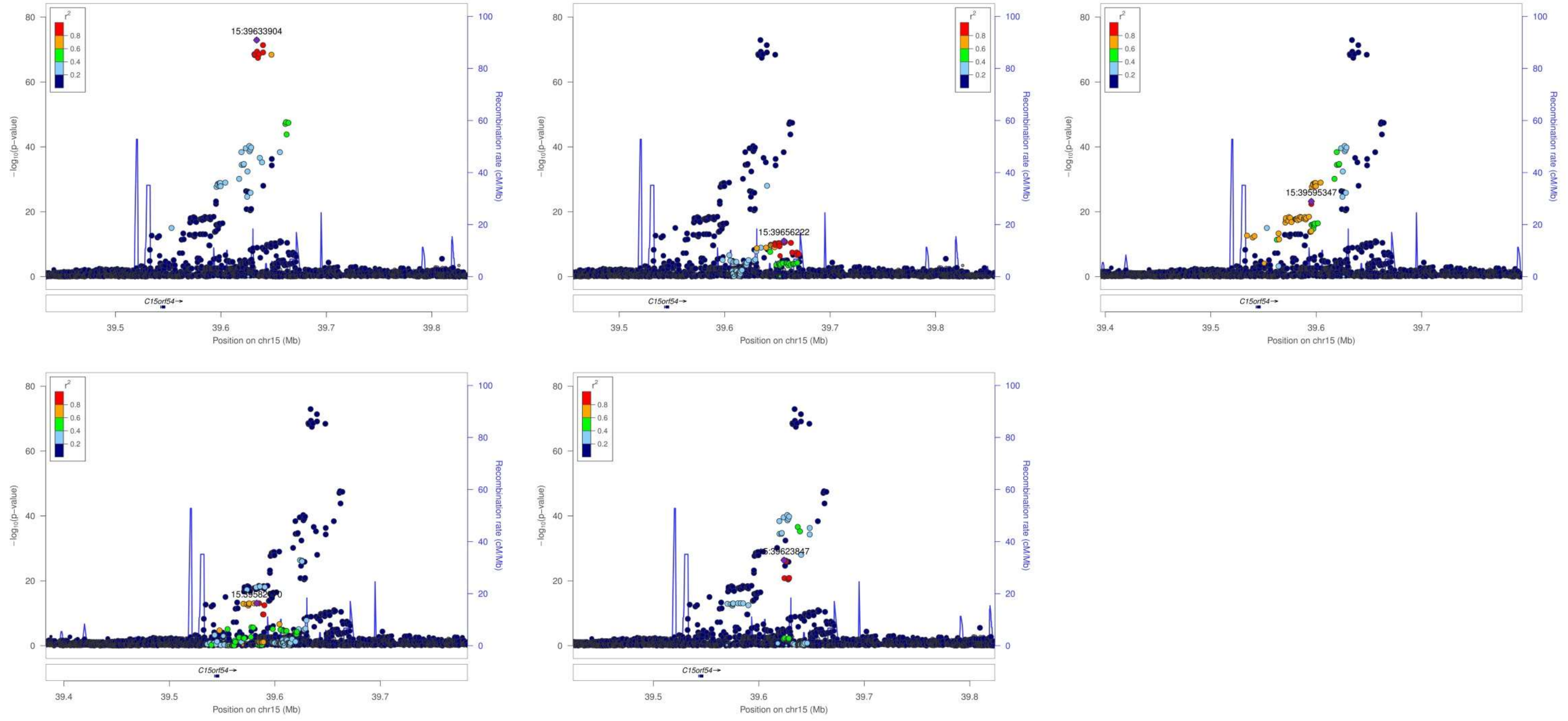

**CSA**

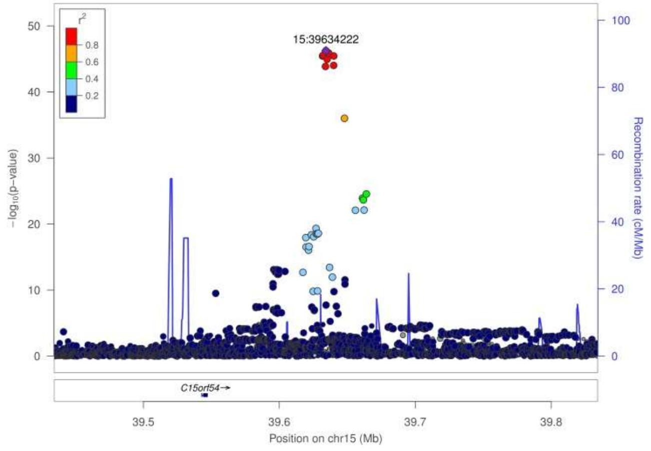
15:39634222
r²
−log10(p−value)
Recombination rate (cM/Mb)
C15orf54→
Position on chr15 (Mb)

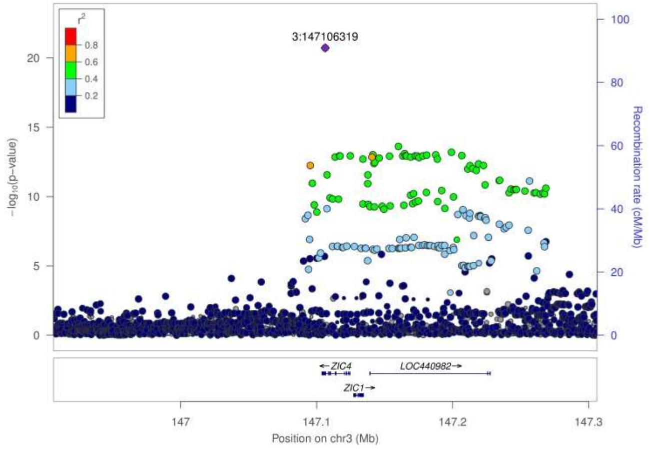
3:147106319
r²
−log10(p−value)
Recombination rate (cM/Mb)
←ZIC4
LOC440982→
ZIC1→
Position on chr3 (Mb)

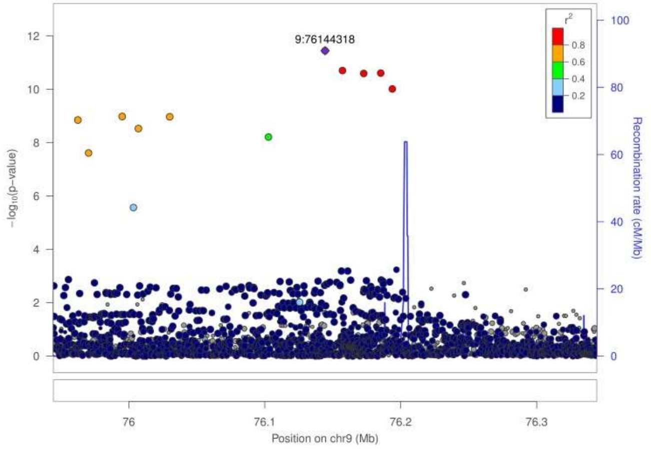
9:76144318
r²
−log10(p−value)
Recombination rate (cM/Mb)
Position on chr9 (Mb)

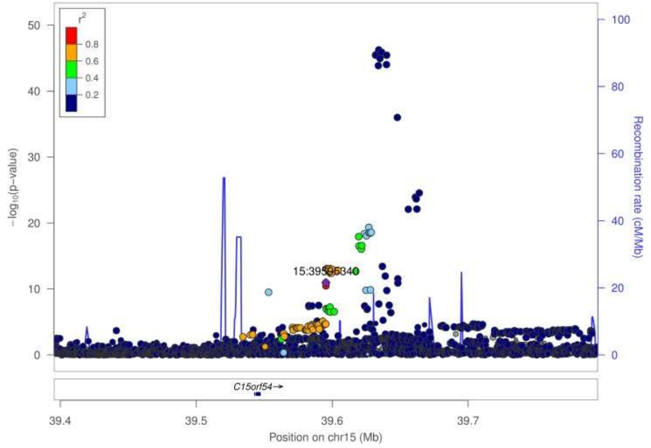
r²
−log10(p−value)
Recombination rate (cM/Mb)
C15orf54→
Position on chr15 (Mb)

CV

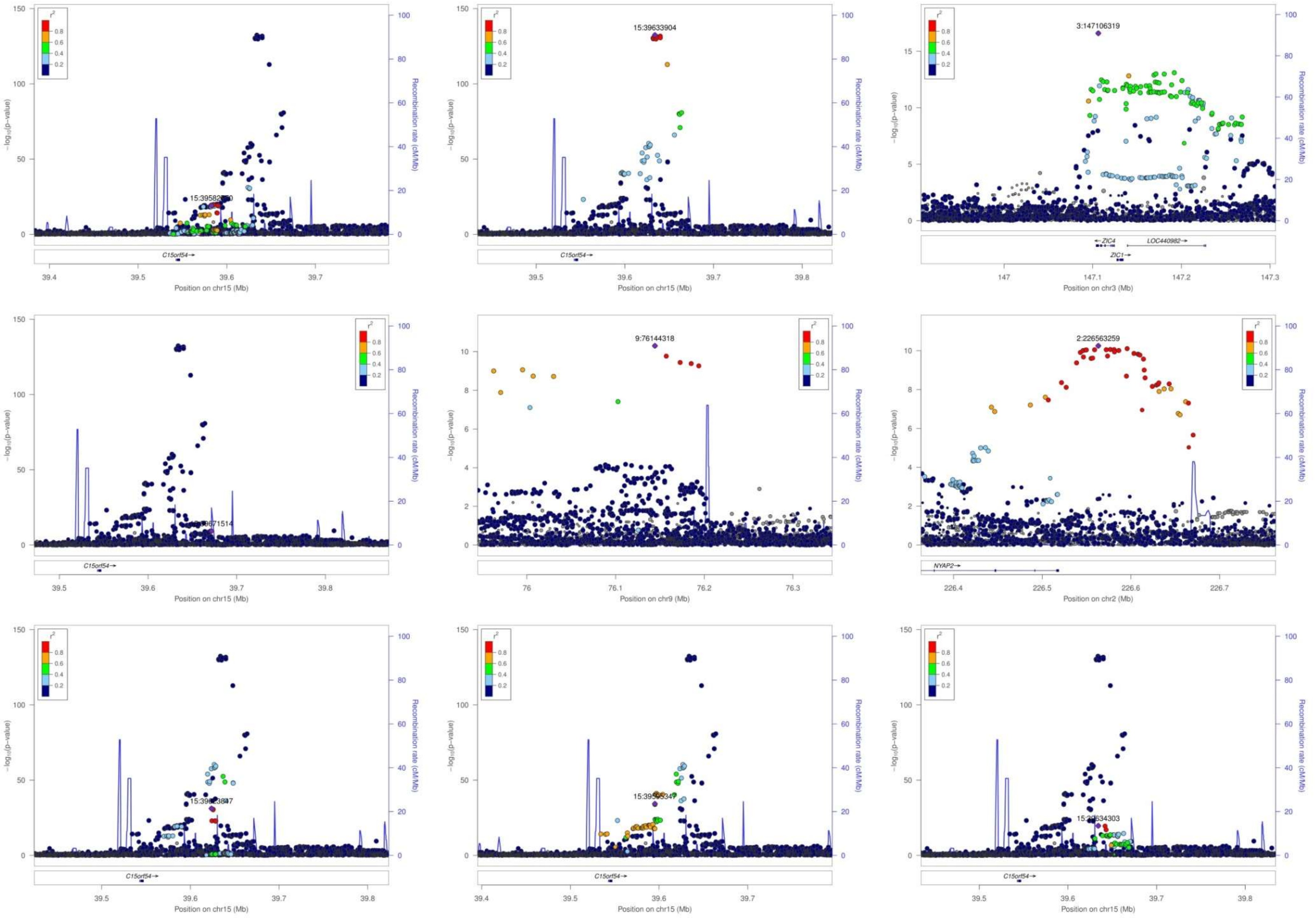

15:39633904
3:147106319
9:76144318
2:226563259
15:39595347
−log10(p−value)
Recombination rate (cM/Mb)
Position on chr15 (Mb)
Position on chr3 (Mb)
Position on chr9 (Mb)
Position on chr2 (Mb)
C15orf54
ZIC4
ZIC1
LOC440982
NYAP2

**Supplementary Figure 4. QQ plots of global cortical thickness (CTh), surface area (CSA) and volume (CV) meta-analyses.**

**global CTh**

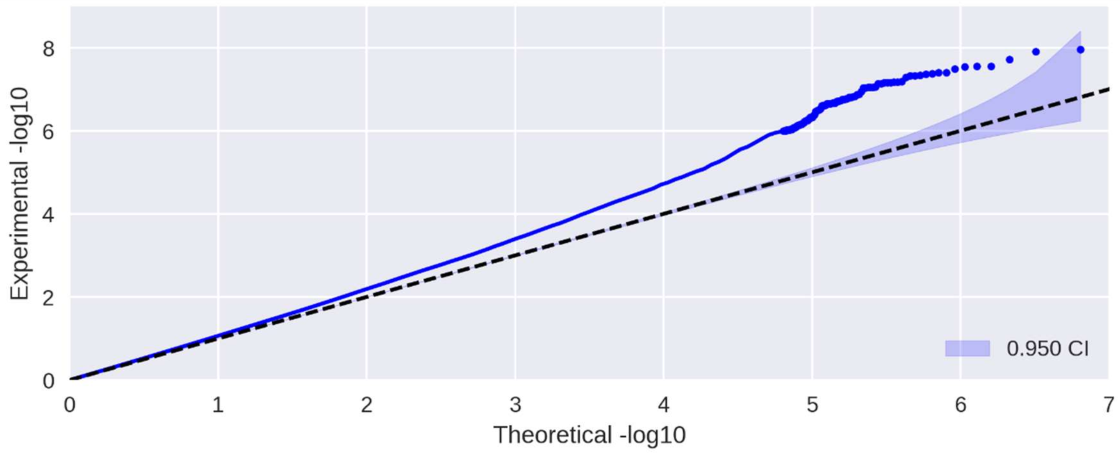


**global CSA**

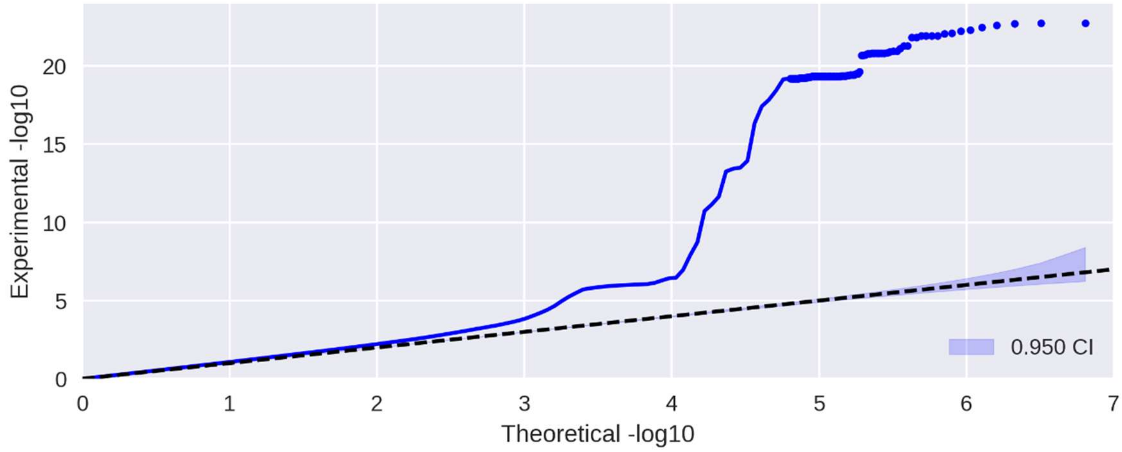


**global CV**

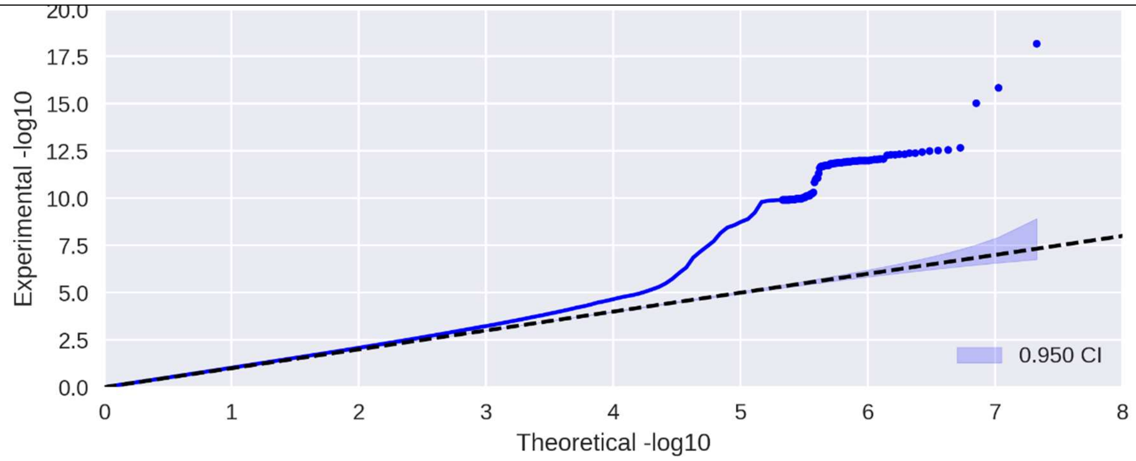

### Supplementary Figure 5. QQ plots of regional cortical thickness meta-analyses.

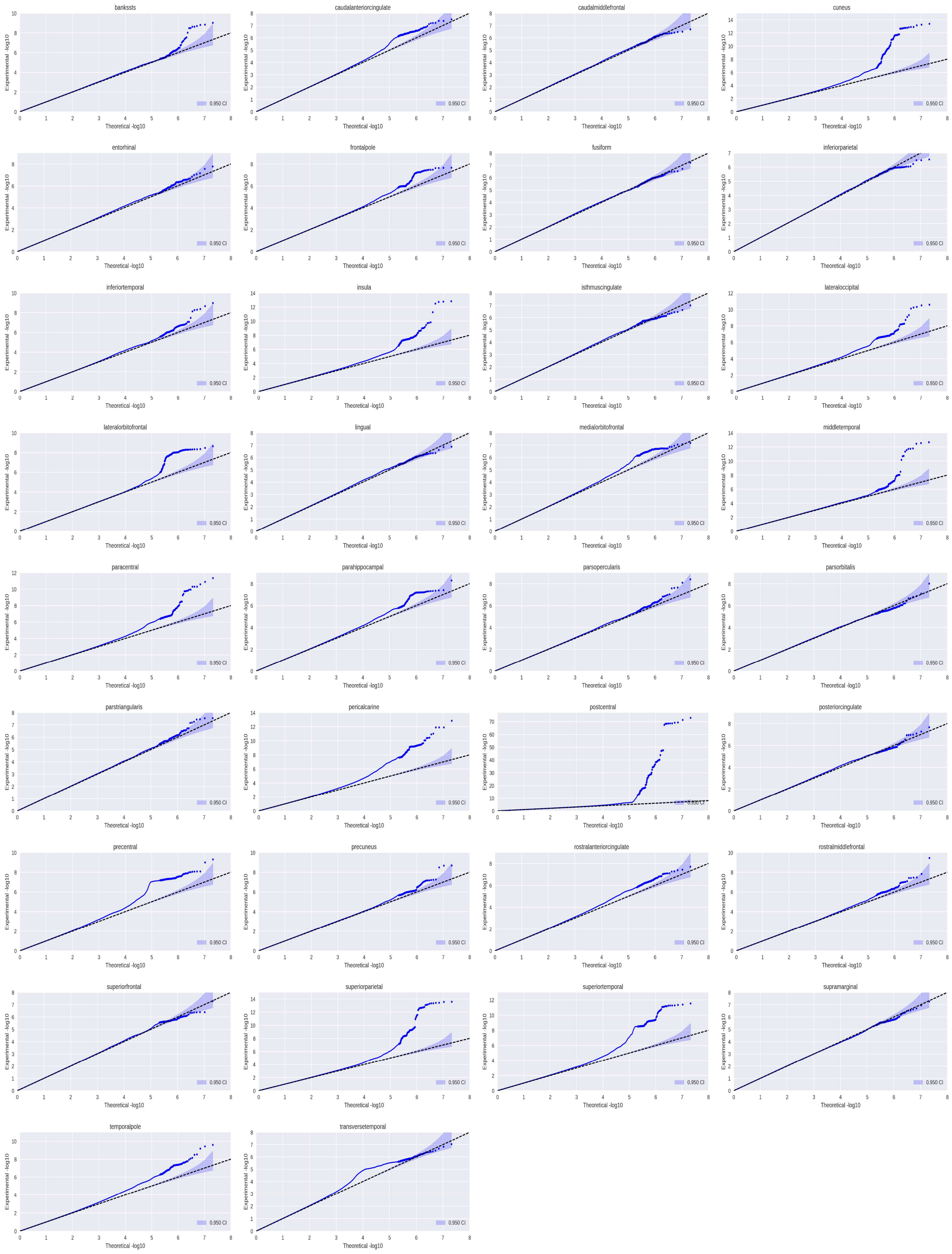

**Supplementary Figure 6. QQ plots of regional cortical surface area meta-analyses.**

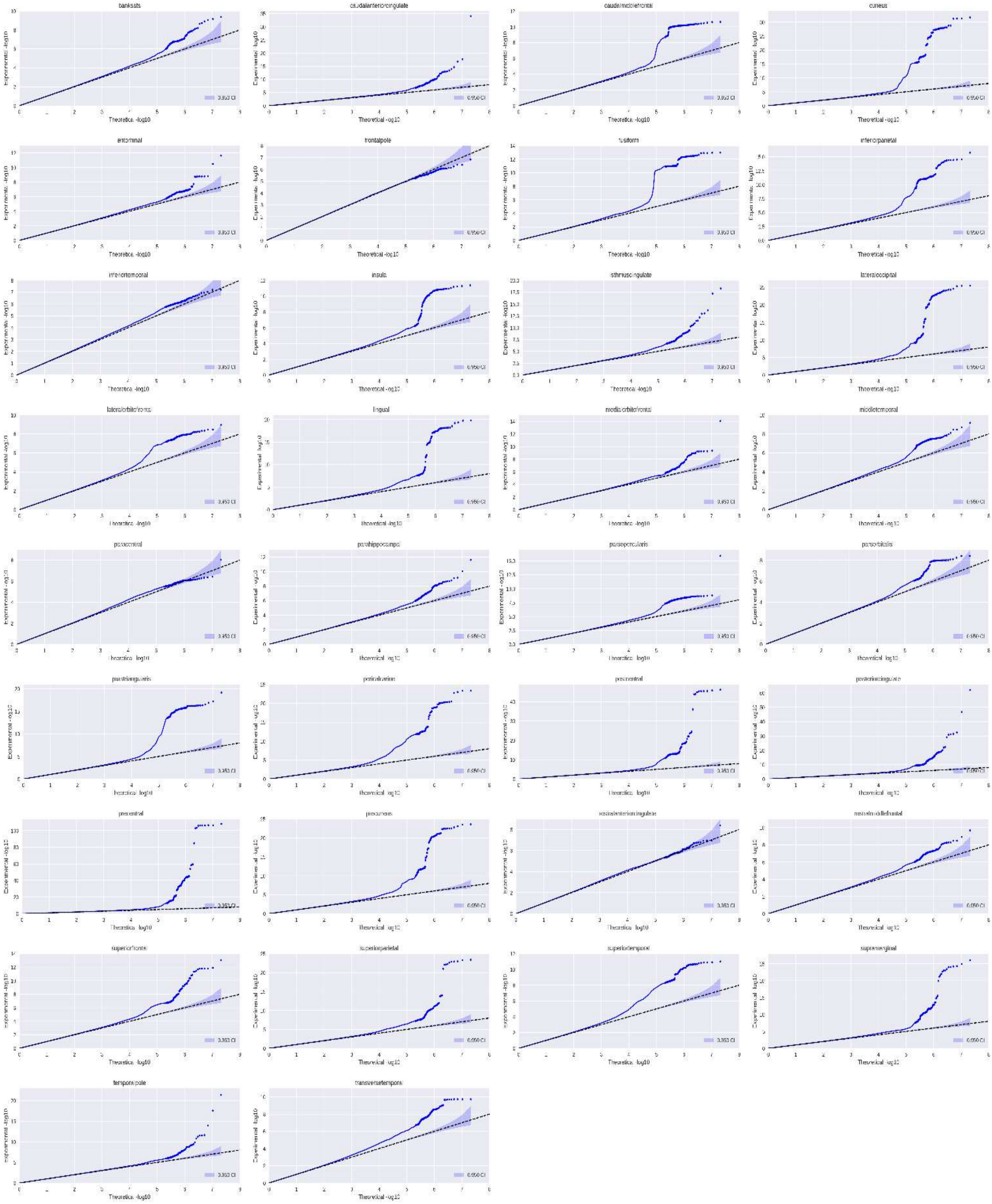

**Supplementary Figure 7. QQ plots of regional cortical volume meta-analyses.**

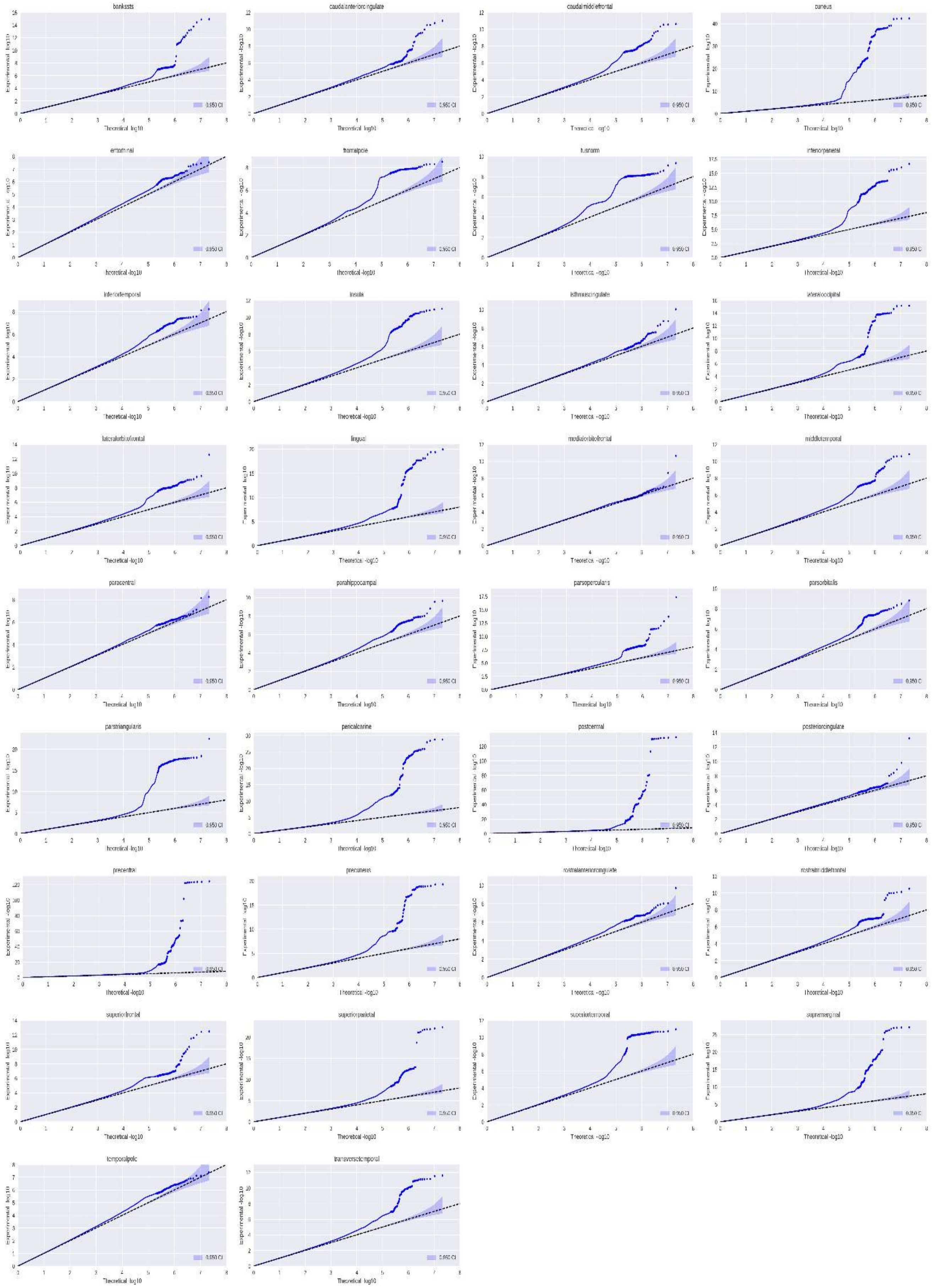

**Supplementary Figure 8. Pathway analysis of the 44 genes that mapped to independent lead SNPs of cortical thickness.**

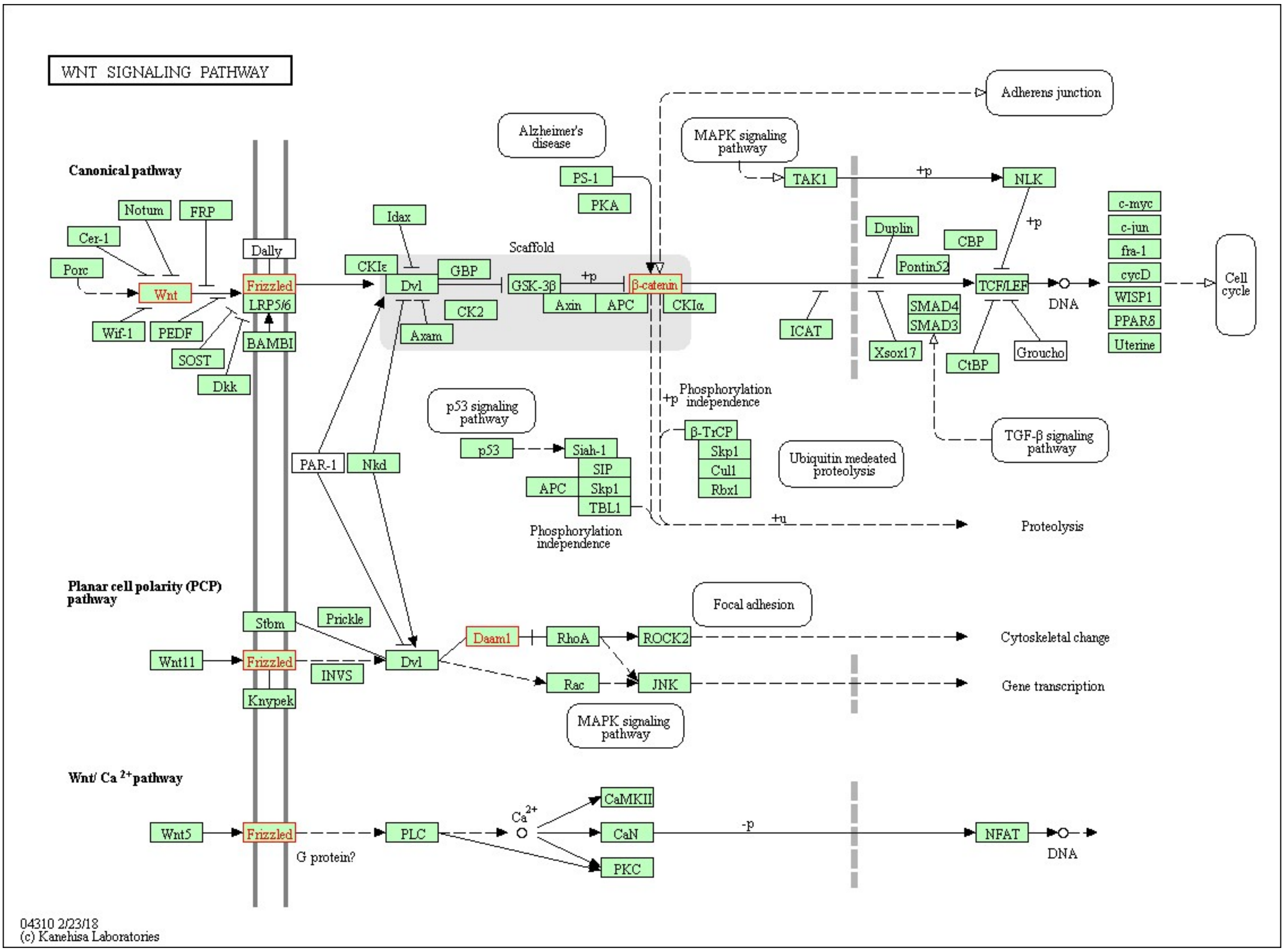


(a)

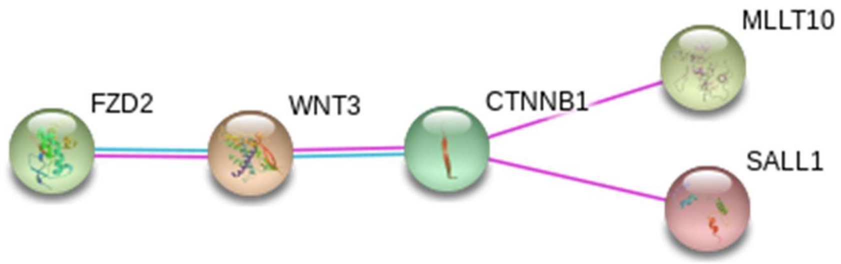


(b)

(a) The Wnt signaling pathway with 140 annotated genes based on KEGG[31] has a significant overlap with those 44 genes that were mapped to independent lead SNPs of cortical thickness (Table S17). The overlap includes *CTNNB1*, *DAAM1*, *DAAM2*, *FZD2*, and *WNT3*, which are shown in red (corrected p-value of hypergeometric test = 0.001).
(b) The protein-protein interactions have been experimentally confirmed by other scholars for three gene products in the overlap. CTNNB1 also interacts with MLLT10 and SALL1, which are among the list of 44 genes with a SNP.

**Supplementary Figure 9. Pathway analysis of the 105 genes that mapped to independent lead SNPs of cortical surface area.**

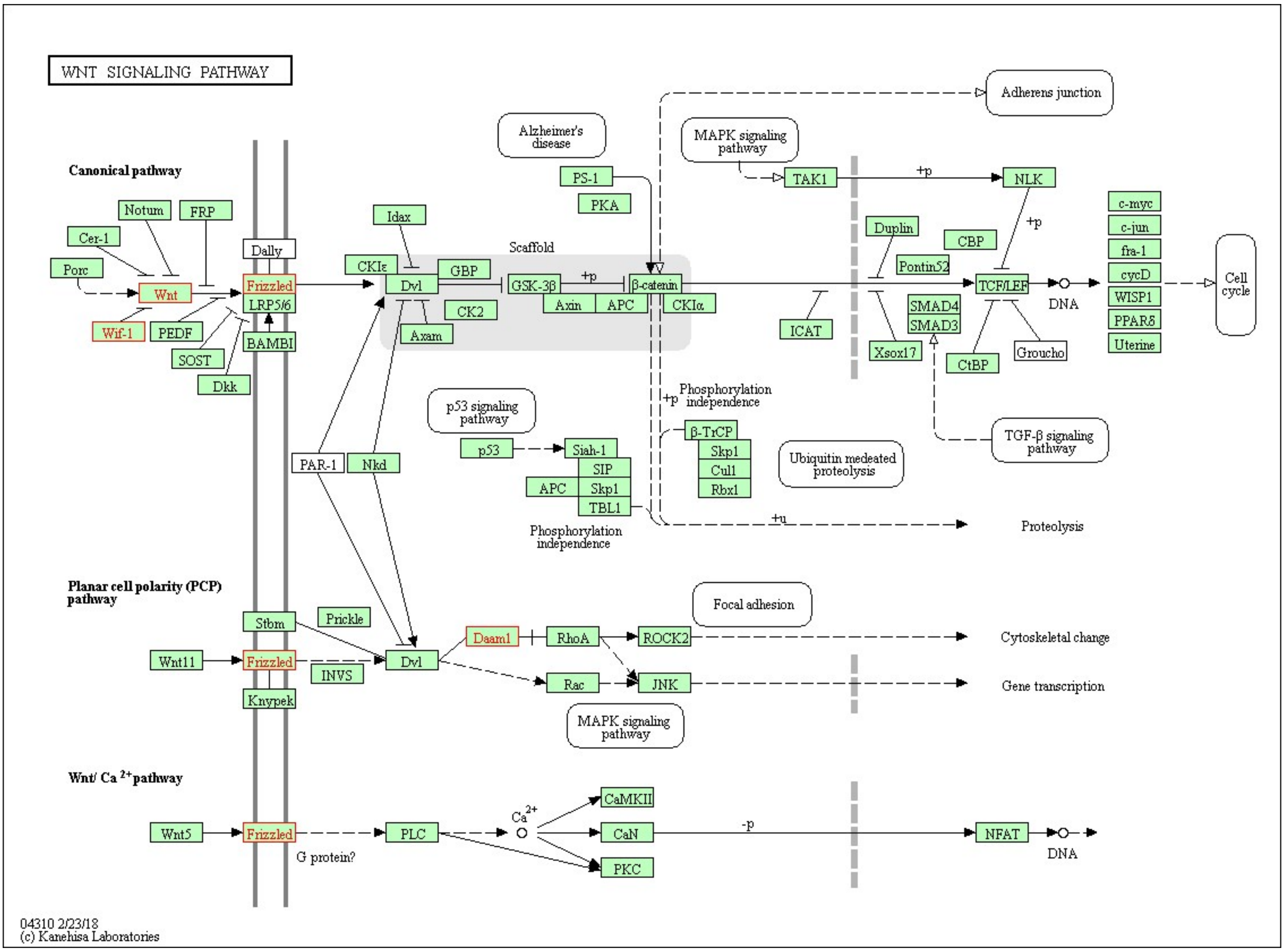


(a)

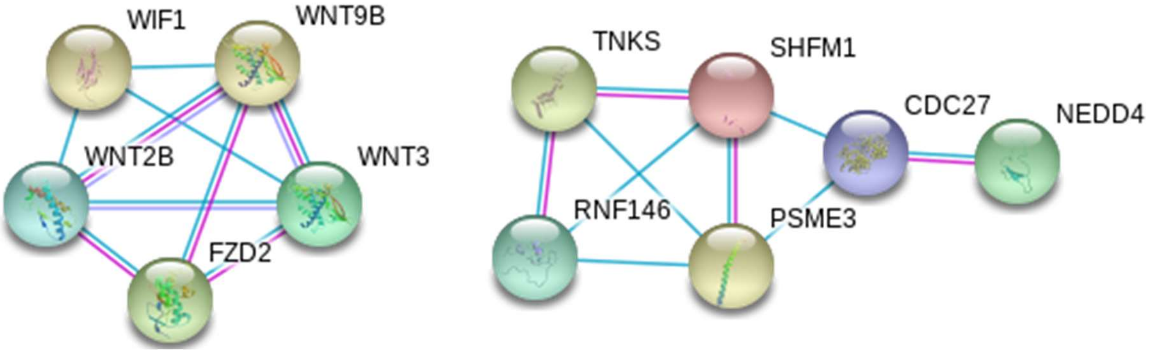


(b)

(a) The Wnt signaling pathway has a significant overlap with those 105 genes that mapped to independent lead SNPs of cortical surface area (Table S18). The overlap includes *DAAM2*, *FZD2*, *WIF1, WNT2B, WNT3,* and *WNT9B*, which are shown in red (corrected p-value of hypergeometric test = 0.005).
(b) The protein-protein interactions relate a subset of these genes in two subnetworks.

**Supplementary Figure 10. Pathway analysis of the 82 genes that mapped to independent lead SNPs of cortical volume.**

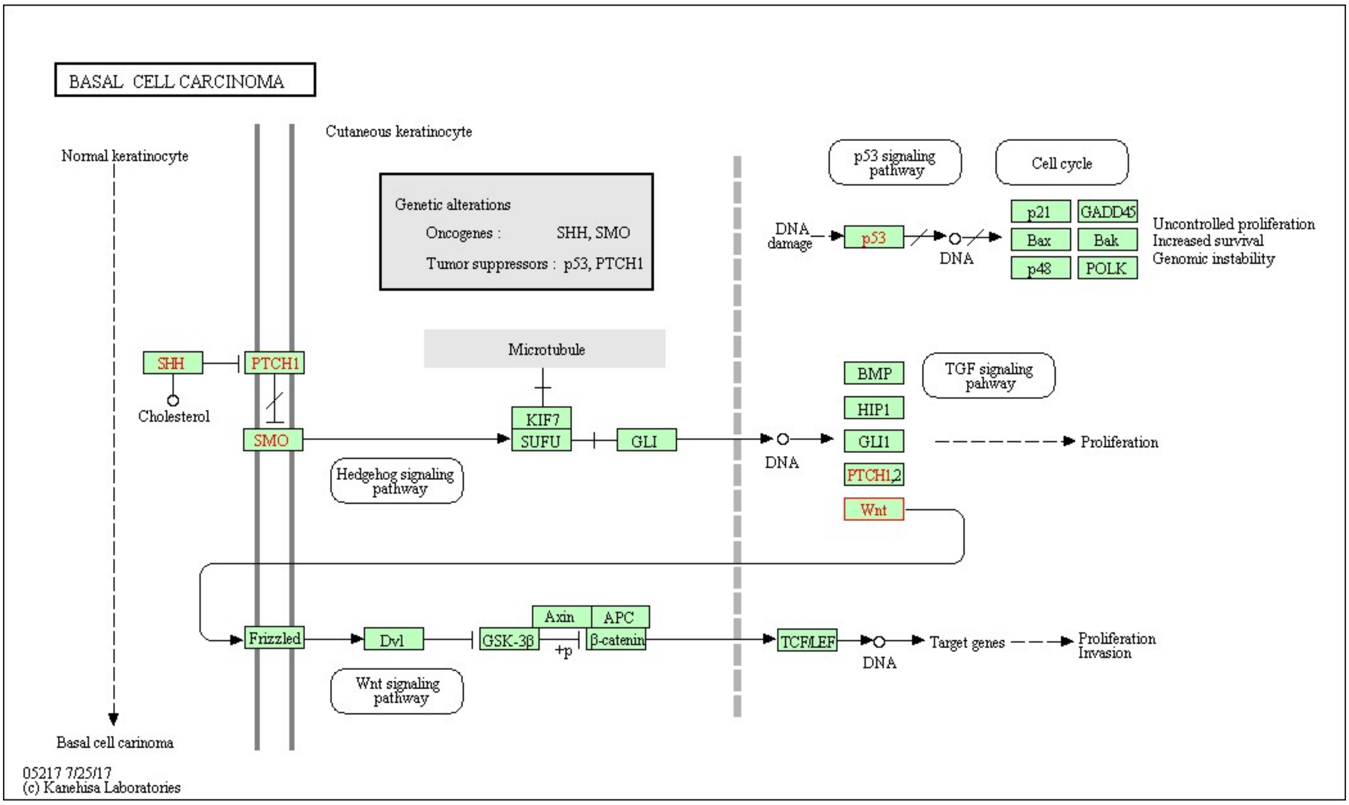


(a)

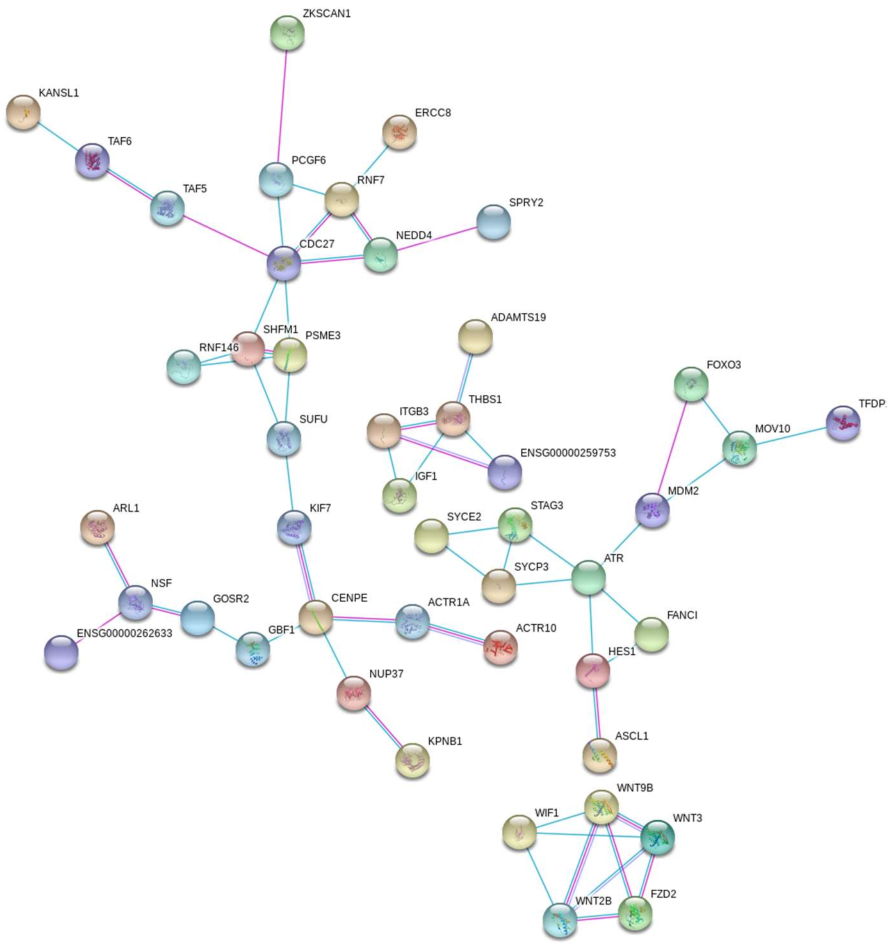


(b)

(a) The 82 genes that mapped to independent lead SNPs of cortical volume (Table S19) have overlaps with the basal cell carcinoma pathway, which has 55 annotated genes based on KEGG. The overlap includes *FZD2, SUFU, WNT2B, WNT3,* and *WNT9B* (corrected p-value of hypergeometric test = 0.04).
(b) Many of these genes have confirmed protein-protein interactions with each other.

**Supplementary Figure 11. Regional heritability estimates based on common SNPs.**

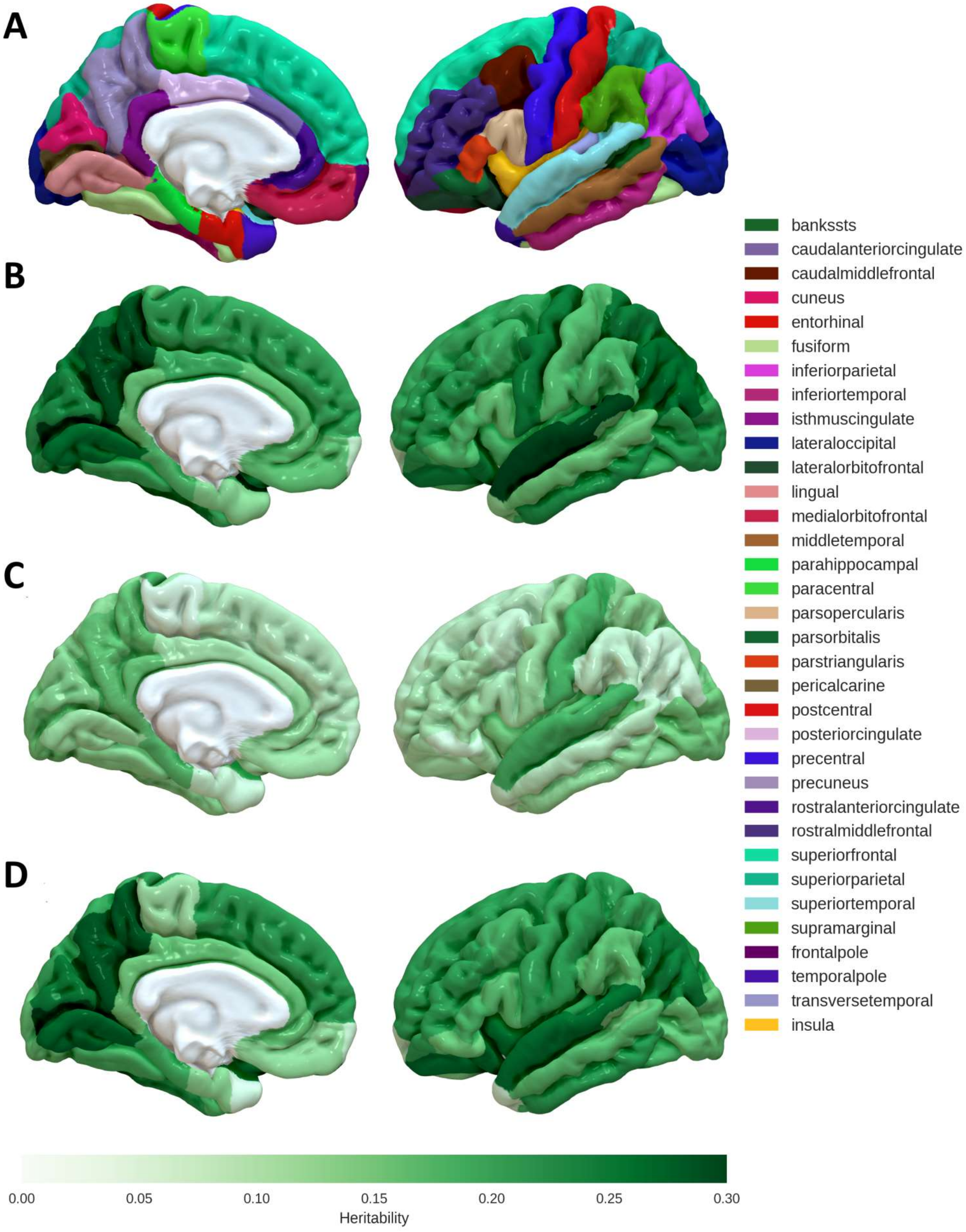


Regional heritability estimates based on common variants, calculated using LD score regression, for (B) cortical surface area, (C) cortical thickness and (D) cortical volume. Panel (A) shows the Freesurfer segmentation.

**Supplementary Figure 12. Genetic correlation between cortical thickness, surface area and volume within cortical regions.**

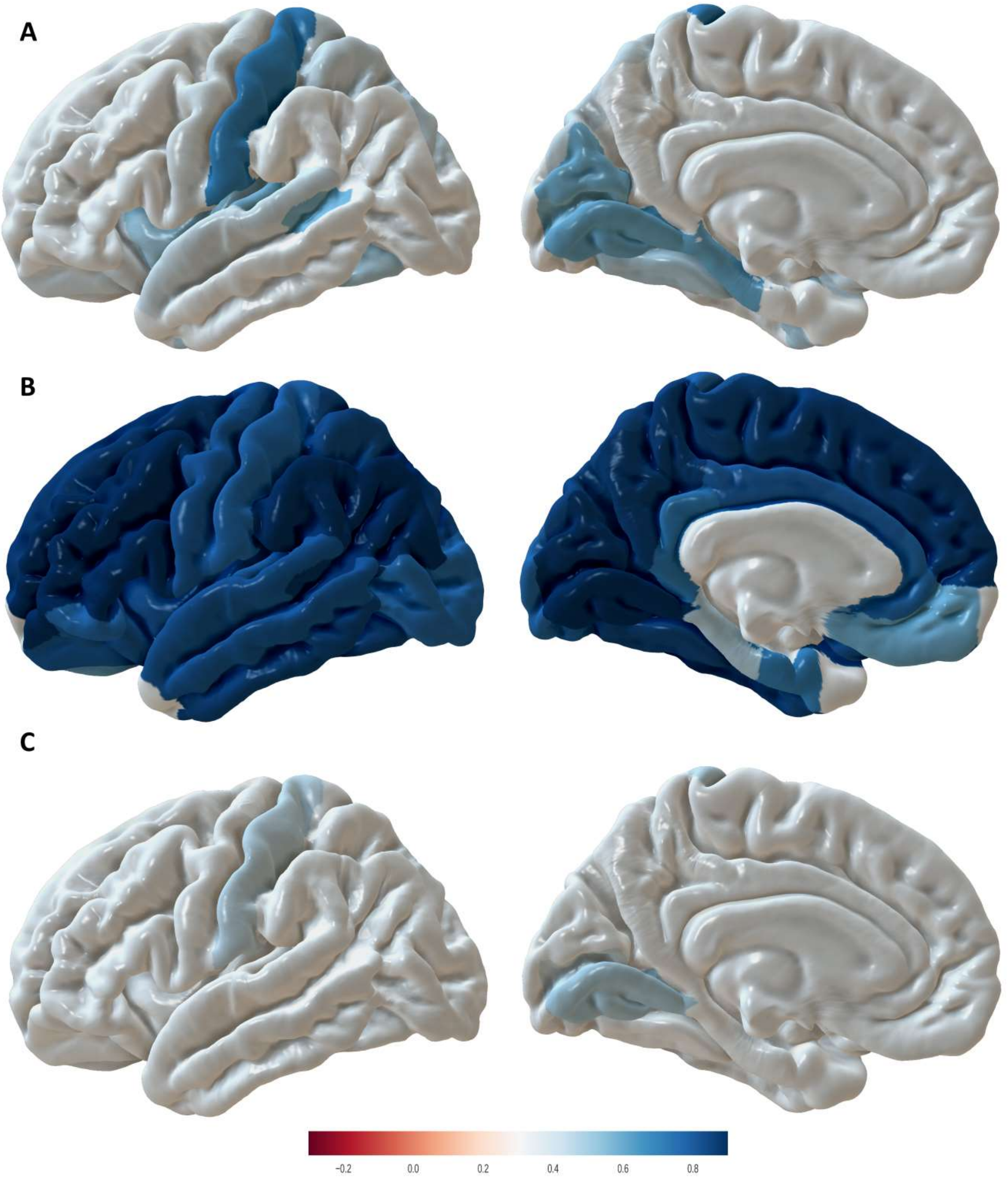


Genetic correlation between cortical thickness and volume (A), cortical volume and surface area (B), and cortical thickness and surface area (C) within cortical regions.

**Supplementary Figure 13. Genetic correlation between regional cortical thickness.**

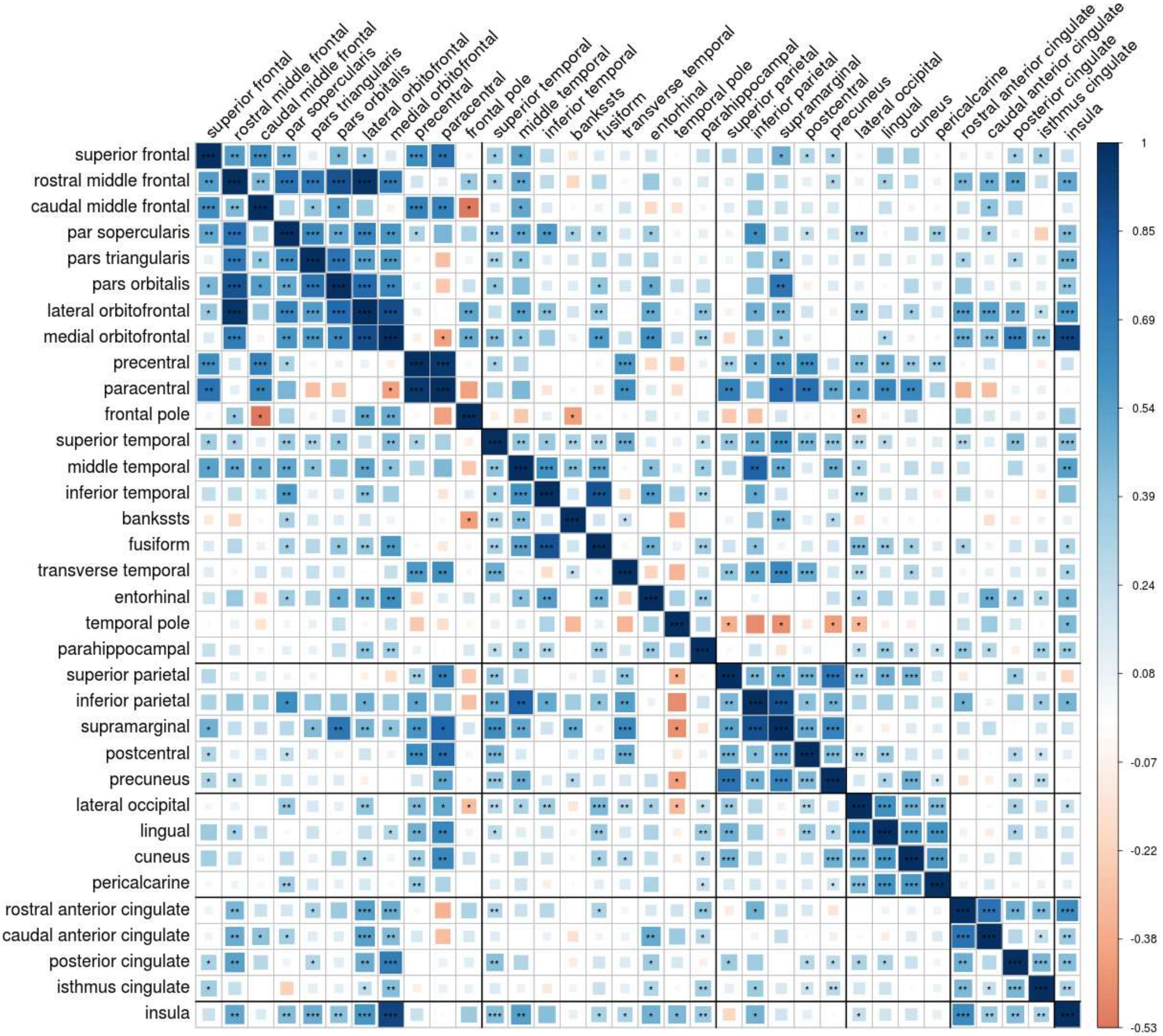


* p-value<0.05, ** p-value<0.01, *** p-value<8.91e-5 (Bonferroni correction: 0.05 (nominal significance) / 34 * (34-1) (number of regions) / 2 (half of the matrix); bankssts=banks of the superior temporal sulcus.

**Supplementary Figure 14. Genetic correlation between regional cortical surface area.**

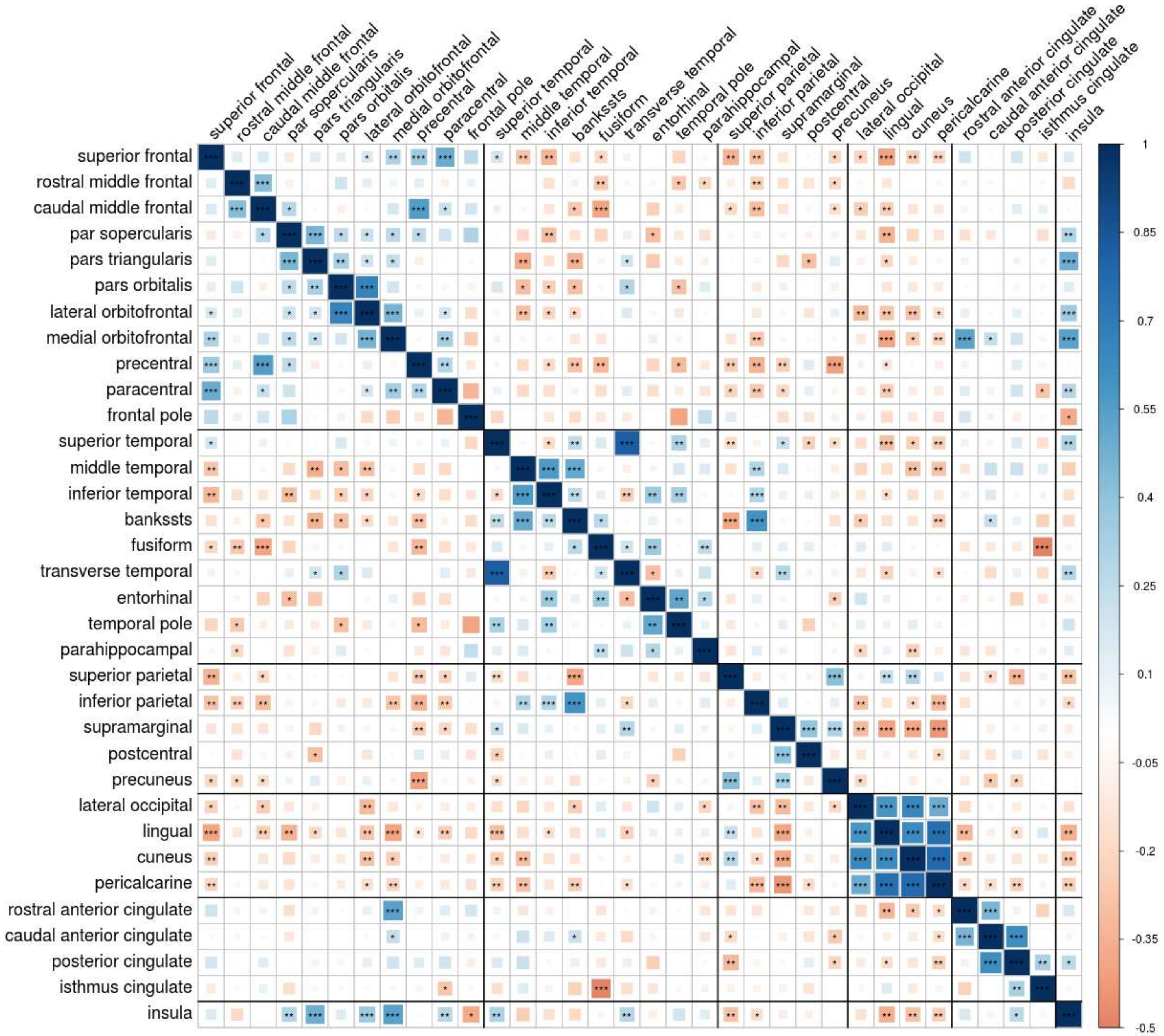


* p-value<0.05, ** p-value<0.01, *** p-value<8.91e-5 (Bonferroni correction: 0.05 (nominal significance) / 34 * (34-1) (number of regions) / 2 (half of the matrix); bankssts=banks of the superior temporal sulcus.

**Supplementary Figure 15. Genetic correlation between regional cortical volume.**

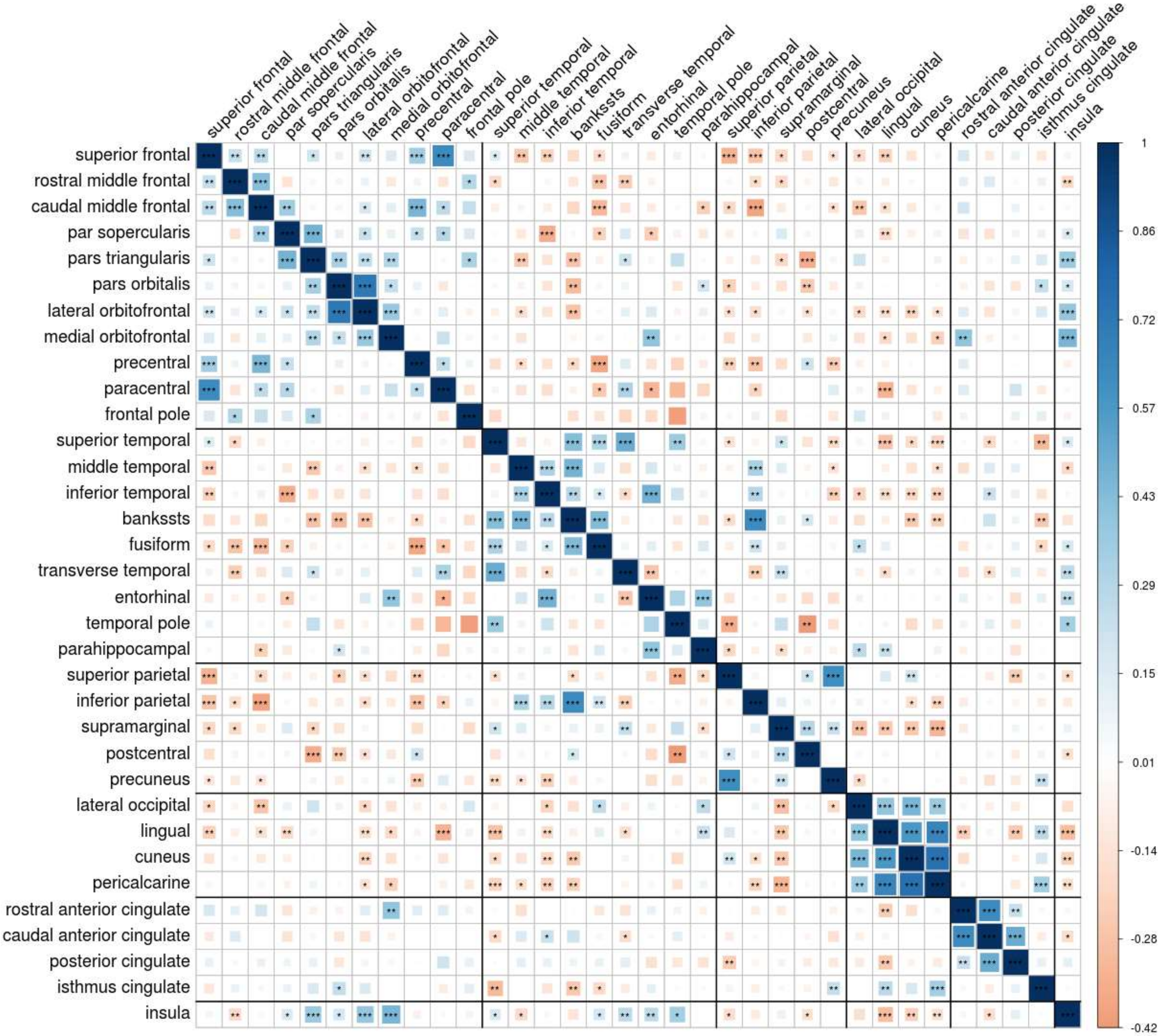


* p-value<0.05, ** p-value<0.01, *** p-value<8.91e-5 (Bonferroni correction: 0.05 (nominal significance) / 34 * (34-1) (number of regions) / 2 (half of the matrix); banksts=banks of the superior temporal sulcus.

**Supplementary Figure 16. Genetic correlation between cortical thickness and other GWAS phenotypes.**

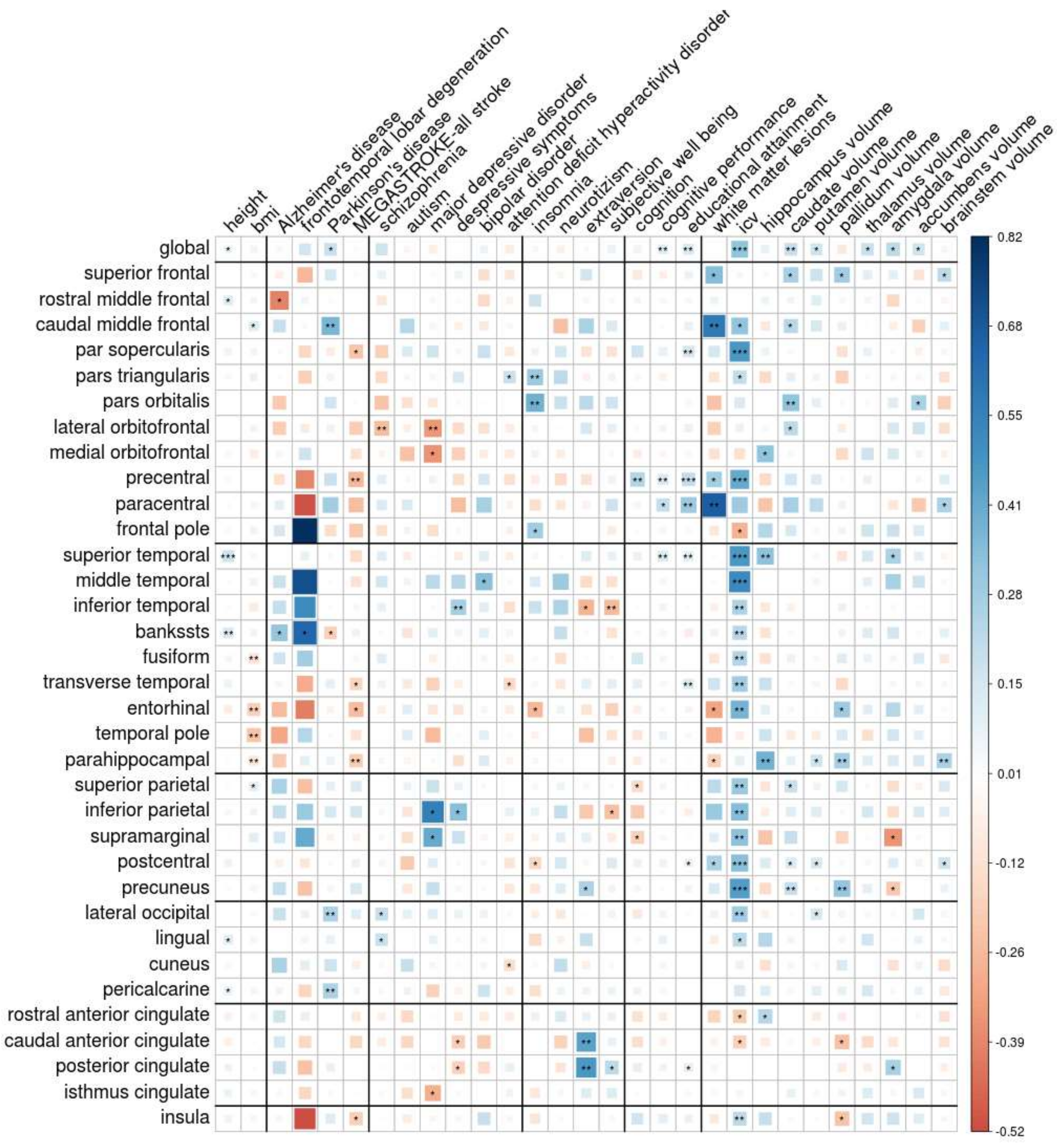


* p-value<0.05, ** p-value<0.01, *** p-value=3.1*10$^{-5}$ (Bonferroni correction: 0.05 (nominal significance) / 46 (number of independent tests)/35(number of tested traits); bankssts=banks of the superior temporal sulcus.

**Supplementary Figure 17. Genetic correlation between cortical surface area and other GWAS phenotypes.**

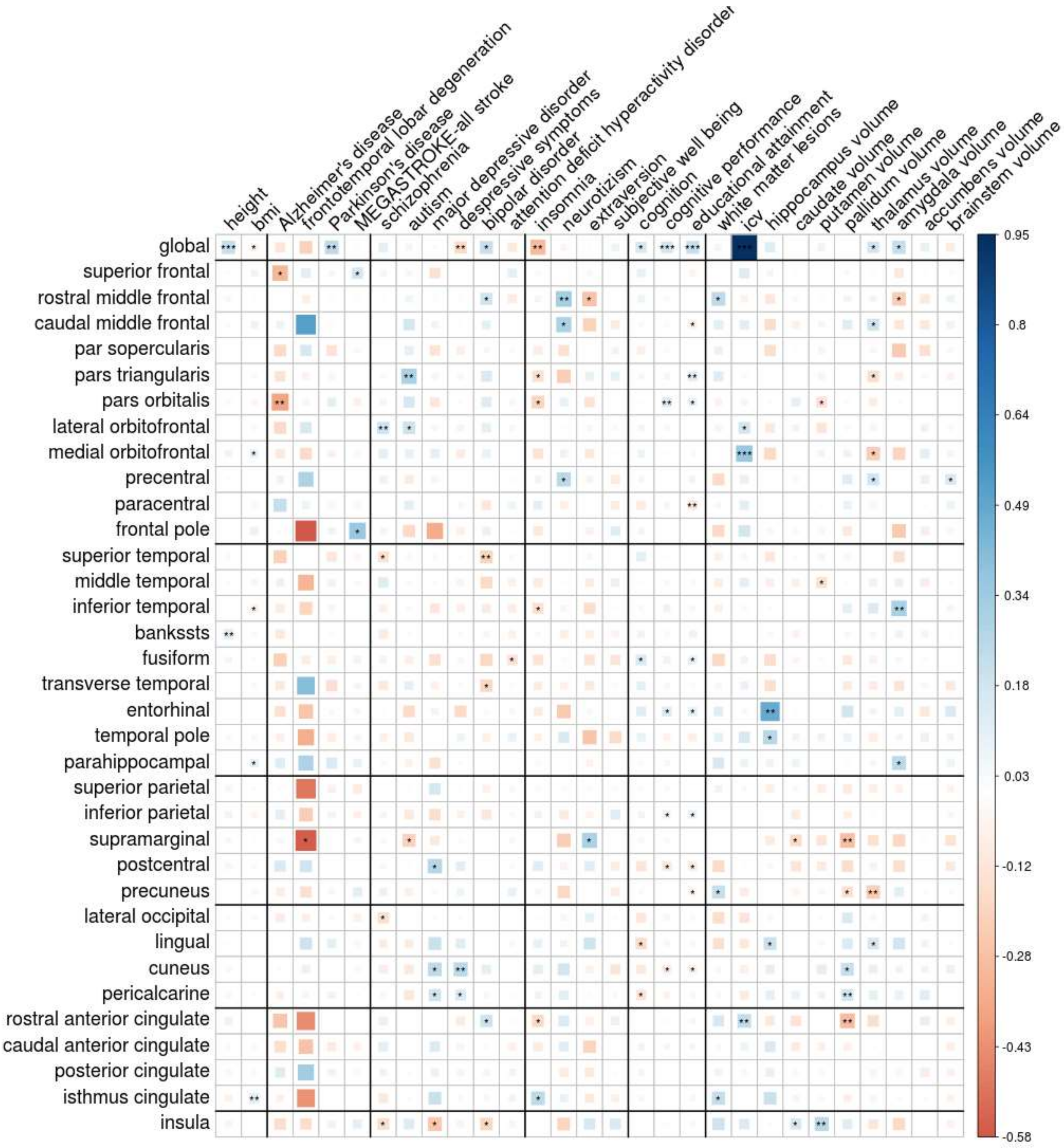


* p-value<0.05, ** p-value<0.01, p-value=3.1*10$^{-5}$ (Bonferroni correction: 0.05 (nominal significance) / 46 (number of independent tests)/35(number of tested traits); banksts=banks of the superior temporal sulcus.

**Supplementary Figure 18. Genetic correlation between cortical volume and other GWAS phenotypes.**

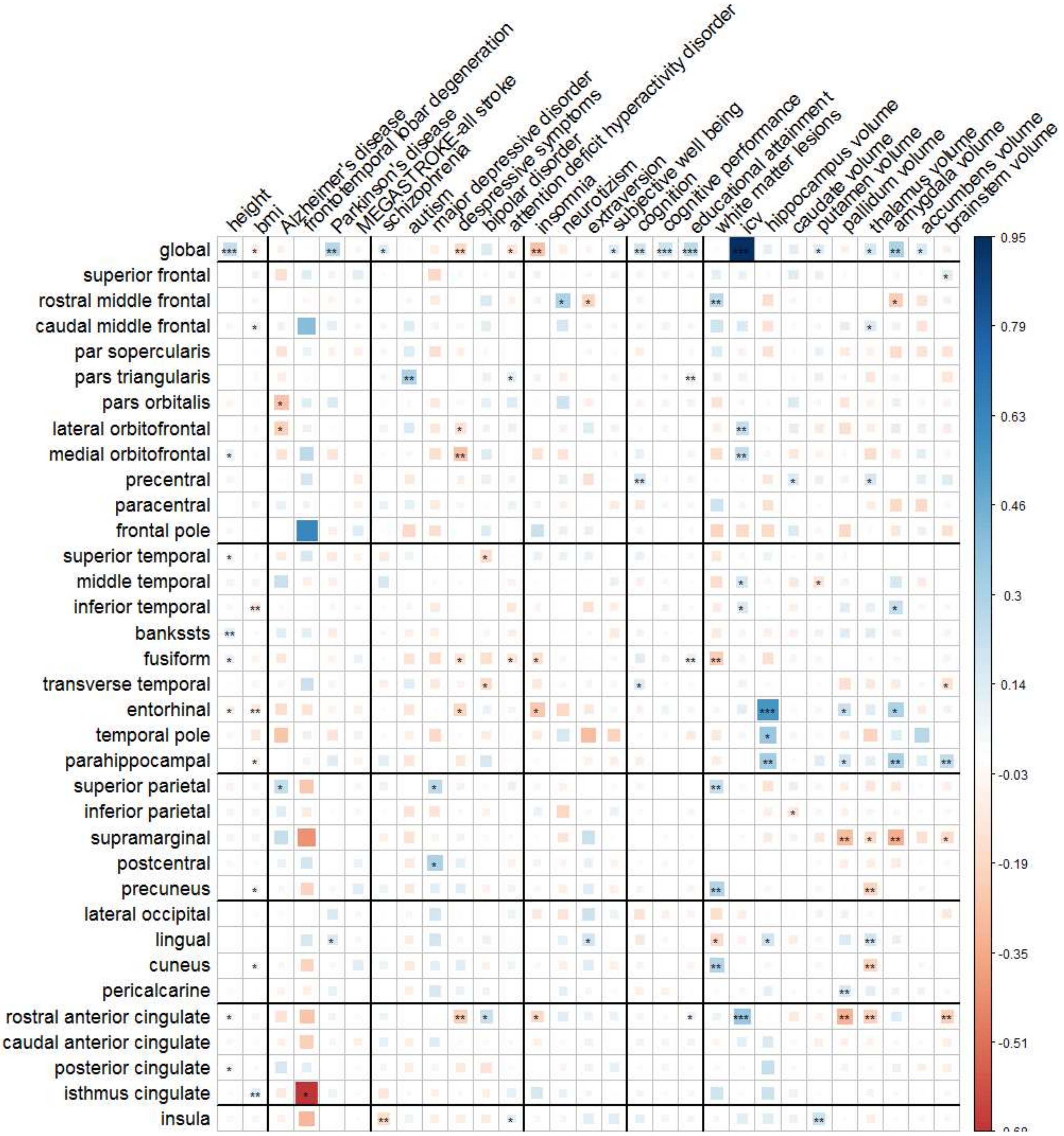


* p-value<0.05, ** p-value<0.01, *** p-value=3.1*10$^{-5}$ (Bonferroni correction: 0.05 (nominal significance) / 46 (number of independent tests)/35(number of tested traits); banksts=banks of the superior temporal sulcus.
